# Insights into the desert living skin microbiome: geography, soil depth, and crust type affect biocrust microbial communities and networks in Mojave Desert, USA

**DOI:** 10.1101/810002

**Authors:** Nuttapon Pombubpa, Nicole Pietrasiak, Paul De Ley, Jason E Stajich

## Abstract

Biocrusts are the living skin of drylands, comprising diverse microbial communities that are essential to desert ecosystems. Although we have extensive knowledge on biocrust ecosystem function and what drives biodiversity in lichen and moss dominated biocrusts, much less is understood about the impacts of diversity among microbial communities. Moreover, most biocrust microbial composition studies have primarily focused on bacteria. We used amplicon-based metabarcode sequencing to explore composition of both fungal and bacterial communities in biocrusts. Specifically we tested how geography, soil depth, and crust type structured biocrust microbial communities or fungal-bacterial networks. Microbial communities were surveyed from biocrust surface and subsurface soils collected from Joshua Tree National Park, Granite Mountain, Kelso Dunes, and Cima volcanic flows located within the Mojave Desert, USA. Five biocrust types were examined: Light-algal, Cyano-lichen, Green-algal lichen, Smooth moss, and Rough moss crust types. We found the primary characteristics structuring biocrust microbial diversity were 1) geography, as central and southern Mojave sites displayed different community signatures, 2) presence of plant associated fungi (plant pathogens and wood saprotrophs), indicator, and endemic species were identified at each site, 3) soil depth patterns, as surface and subsurface microbial communities were distinctly structured, and 4) the crust type, which predicted distinct microbial compositions. Network analysis showed that Cyanobacteria and Dothideomycetes (Pleosporales) were the major hubs of overall biocrust microbial community. Such hierarchical spatial organization of biocrust communities and their associated biotic networks can have pronounced impacts to ecosystem functions. Our findings provide crucial insights for dryland restoration and sustainable management.

## 1. Introduction

In vegetation sparse drylands, plant interspaces are often covered by biological soil crusts (hereafter biocrusts)[1]. Microbial communities form biocrusts by establishing at the uppermost few millimeters of the soil, as interwoven biofilaments forming a living soil aggregate. As a major component of dryland landscapes, biocrusts account for 12% of Earth’s terrestrial surfaces [2,3]. They can be thought of as a living skin of dryland soils embodying hot spots of biodiversity, biogeochemistry, and ecosystem functions. Evolutionarily diverse organisms such as bryophytes, lichens, eukaryotic algae, cyanobacteria, bacteria, and fungi combine to form different types of biocrust distinguished by their dominant photoautotrophic community member [1,3-5]. The complex combinations of microorganisms in biocrusts affect a range of ecosystem functions, such as: mediating soil nutrient cycles, preventing soil erosion and improving soil stabilization, assisting with regeneration of vegetation, as well as fertilizing and transforming subsurface soils [1,2,5,6].

While there is an extensive body of literature on biocrust lichen and bryophyte diversity, as well as their roles in dryland ecosystems, studies have only recently begun exploring their less conspicuous community members. Initial surveys of biocrusts’ microbial composition targeted bacteria and algae [7-10]. As the major photosynthetic components of biocrust, some studies specifically examined cyanobacteria and eukaryotic algae, but the majority of these focused on European biocrusts [11-15]. Knowledge from other continents is as yet much less extensive [16]. Most published work has employed culture dependent approaches, which likely underestimate microbial diversity (as demonstrated by for example ref. 17,18). Few have investigated how photoautotrophic diversity is associated with heterotrophic archaea, bacteria, and fungi in biocrusts. Moreover, microbial community associations were not usually explored in different biocrust types identifiable by their dominant photoautotrophic community member as light (cyanobacterial/algal), dark (cyanobacterial/algal), lichen, and bryophyte crust [3,19,20].

In particular, our knowledge of biocrust fungal diversity is extremely poor. In the past, biocrust fungal diversity was examined using denaturing gradient gel electrophoresis (DGGE) [21]. Newer procedures greatly improve biodiversity assessment, but only a handful of researchers have applied environmental DNA-based next generation sequencing (NGS) approaches to biocrust systems [3,22,23]. Although general trends of fungal diversity agree between DGGE and NGS, the latter approach offers much greater taxonomic coverage and resolution than DGGE. Specifically when using ITS amplicons, a broad range of fungal diversity can be revealed and be used to identify most taxa to genus or species level [24-26]. In addition to ITS NGS, 16S NGS has been used successfully to characterize archaeal and bacterial diversity [27]. As a result, broad coverage of bacteria, archaea, and fungal diversity in biocrust can now be attained with great taxonomic resolution. Moreover, NGS of combined microbial taxa allows us to answer microbial linkage questions which have not yet been explored in biocrust communities. Thus, joint surveying of fungal and bacterial/archaean communities with NGS data enables deeper comparisons of biocrust microbiome diversity, allowing us not only to identify core taxonomic composition of biocrust types, but also to specify the key players and thus reveal network connections within bacterial communities, within fungal communities, and also across fungal-bacterial communities.

Similarly less explored are questions about regional patterning of biocrust diversity, or how diversity changes vertically when comparing the biocrust to the adjoining subsurface soil. When microbial communities were studied with DGGE, similar major fungal and bacterial phyla were observed in biocrusts from different localities [9,28,29]. However, sampling was often conducted at a small spatial extent, and the procedures employed often focused on a single group of organisms. Although similar major phyla were found in different localities, previous results suggested different relative abundance geographically. To date, only a handful of studies reported broadscale geographical patterns of biocrust microbial communities in North America [30-32]. Moreover, most biocrust studies focused on the biocrust itself (surface) without comparing the underlying subsurface microbial community. Yet recent investigations have shown differences between the surface and subsurface bacterial community [8,28,33]. Such vertical heterogeneity may create confounding effects when comparing alpha/beta diversity results from different surveys especially if standard soil depths are used in sampling efforts. Recognizing these gaps in our understanding gives us the opportunity to investigate microbial community composition and structure at different levels of complexity - regionally, structurally among biocrust types, as well as vertically by soil depth. Therefore, a regional survey of biocrust microbial community survey is needed which also focuses on microscale patterns within each sampling site by incorporating different crust types and soil depths.

We investigated microorganisms from the three domains of life including Archaea, Bacteria, and Fungi, using high-throughput amplicon sequencing targeting both the 16S rRNA and ITS1 markers. In this paper we ask the following research questions: 1) Does geographical location structure biocrust microbial communities? 2) Do biocrusts contain unique (endemic) microbial species at each site? 3) Do biocrust microbial surface communities differ from those in the adjacent soil subsurface? 4) Do biocrust types each have their own unique assemblages of microbes and do characteristic differences in richness exist between them?

To address these questions, we collected biocrust samples from four different sites along a north-south axis within the Mojave and the ecotone of the Mojave and Colorado Deserts. We collected both surface and subsurface material from five different biocrust types from each site. We hypothesized that: 1) geographical locations structure biocrust microbial communities where our 3 central Mojave sites will have similar microbial composition while the JTNP site at the ecotone of the Mojave and Colorado desert will have different microbial composition; 2) different geographical locations will harbor key/endemic species that are unique to each site; 3) soil depth affects fungal and bacterial diversity in which light dependent microbes have higher abundances on the surface while heterotrophs are more abundant below the biocrust. Both alpha and beta diversity will distinguish subsurface soil microbial community composition from surface communities, where photoautotrophs are major components; and 4) biocrust types relate to microbial diversity, where more complex assemblages such as lichen and moss crusts will have greater alpha diversity in both fungal and bacterial composition than structurally less complex types such as light algal crusts.

## 2. Materials and Method

### 2.1 Sampling sites and biocrust sampling

Biocrust samples were collected from four different sites in the Mojave Desert and at its southern edge. Our Joshua Tree National Park (JTNP, GPS: 34.10N, -115.45W) site is located at the ecotone of the Mojave Desert with the Colorado Desert, while sites at Granite Mountains (GMT, GPS: 34.78N, -115.63W), Kelso Dunes (KELSO, GPS: 34.89N, -115.69 W), and Cima volcanic field (CIMA, GPS: 35.20N, -115.87W) sites were located further north in the central Mojave Desert (Figure 1D) (Google, 2019). Using sterile sampling technique, five biocrust types were collected with a spatula. The underlying subsurface soil for each biocrust type was also collected (Figure 1G) by pushing a 5cm diameter brass core to a depth of 5cm (or less if subsurface rock interfered). Light algal crust (LAC, Figure 1A), Cyanobacteria lichen crust (CLC, *Collema* spp., Figure 1C) and Green algal lichen crust (GLC, *Clavascidium lacinulatum*, Figure 1B & 1E) were collected at all four sites, while rough moss crust (RMC, *Syntrichia* spp., Figure 1F) and smooth moss crust (SMC, *Bryum* spp., Figure 1H) were collected in addition at KELSO, GMT, and CIMA (neither type was sufficiently prevalent for collection at the JTNP sampling site). For each type of biocrust, surface versus subsurface soil samples were collected into separate 50 ml tubes. Biocrust samples were stored on ice and transferred to a -80°C freezer at University of California Riverside.

**Figure 1.**
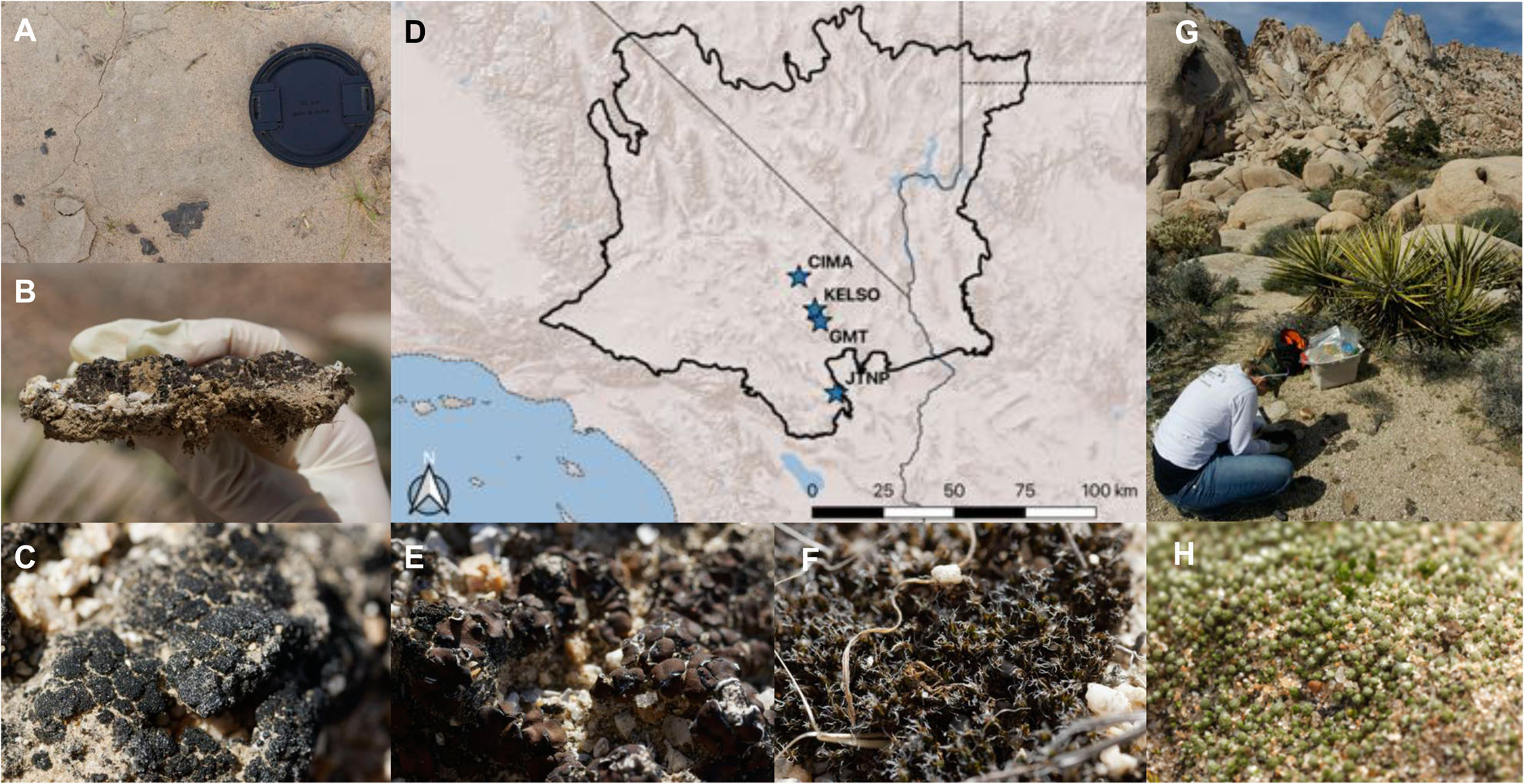
Sampling sites and biocrust types. A) Light algal crust (LAC), B) dangling filaments underneath GLC, C) Cyanobacteria lichen crust (CLC), D) our 4 sampling sites including Cima Volcanic Flows (CIMA), Kelso Sand Dunes (KELSO), Granite Mountains Research Center (GMT) within the Mojave Desert (black outlined area), and Joshua Tree National Park (JTNP) at the edge of the Mojave and Colorado Deserts [33,34], E) Green algal lichen crust (GLC), F) Rough moss crust (RMC), G) crust sampling in Mojave Desert, H) Smooth moss crust (SMC)

### 2.2 NGS sequencing analysis

DNA extraction was performed with 0.15 g of biocrust using the QIAGEN DNeasy PowerSoil kit (Qiagen, Germantown, MD, USA) following manufacture standard protocol. The ITS1F and ITS2 primer pair was used to amplify the internal transcribed spacer region (ITS) for the fungal community according to Smith and Peay’s Illumina MiSeq protocol [24]. The 515F and 806R primers were used to amplify the 16S rRNA V4 gene region for bacterial community following Caporaso et al. [27]. PCR reactions were processed in 25 ul total volume in three replicates which included 1 ul of each primer (10 uM), 1 ul of genomic DNA, 12.5 ul of Taq 2X DNA Polymerase (Thermo Fisher Scientific Inc., Waltham, MA, USA), and 9.5 ul of nuclease-free water (Sigma-Aldrich, St.Louis, MO, USA). PCR conditions were: initial denaturation at 93°C for 3 min; 35 cycles of denaturation at 95°C for 45 sec, annealing at 50°C for 1 min, extension at 72°C for 90 sec, and a final extension at 72°C for 10 min using C1000 thermal cycler (BioRad, Hercules, CA, USA). PCR products from three replicates were combined and then purified using NucleoSpin Gel and PCR Clean-up kit (Macherey-Nagel, Hoerdt, France) and pooled to produce equimolar mixture. Pooled libraries were quantified using Qubit dsDNA HS Assay (Life Technologies, Carlsbad, CA, USA) and analyzed using Agilent 2100 Bioanalyzer and Fragment Analyzer (Agilent Technologies, Santa Clara, CA, USA). Then pooled libraries were sequenced using Illumina MiSeq (San Diego, CA) with the V3 kit to generate paired-end reads in 2 x 300bp format, at the Institute for Integrative Genome Biology, Core Facilities, University of California, Riverside (http://iigb.ucr.edu). A total of 8,918,345 paired end sequence reads were produced and submitted to the Sequence Read Archive (SRA) databases associated with BioProject accession number PRJNA544067.

### 2.3 Bioinformatics

The fungal ITS1 amplicon sequences were analyzed with AMPtk: the Amplicon Toolkit for NGS data (formally UFITS) [34] (https://github.com/nextgenusfs/amptk). The demultiplexed paired-end sequences data were pre-processed by trimming forward and reverse reads to a maximum of 300 bp, trimming primer sequences and discarding reads less than 100 bp in length. The paired-end reads were merged to produce a single long read using USEARCH v9.1.13 [35] where they could be found to overlap. After pre-processing, a total of 3,040,944 valid paired sequence reads were produced. Sequence quality filtering was performed with the expected error parameter of 0.9 [36], which produced 2,392,561 quality filtered reads. This cleaned sequenced dataset was clustered with UPARSE using a 97% percent identity parameter, which generated 2,569 Operational Taxonomic Units (OTUs) following the procedure of Palmer et al. [34]. Chimeric OTUs, sequences produced from PCR amplification of templates or parent sequences, were filtered using VSEARCH (v 2.3.2) [37] which removed 65 chimeras after comparison to the database. Finally, taxonomic assignment for 2,504 OTUs was performed with the AMPtk hybrid approach using names from UNITE v8.0 [38] and functional guilds were assigned using FUNGuild v1.0 [25].

The 16S V4 amplicon sequences were analyzed using Quantitative Insights Into Microbial Ecology version 2 (QIIME2 v2019.1)[39] using bacterial 16S processing workflows. Demultiplexed sequence data (5,757,892 reads) were imported to QIIME2 then pre-processed by trimming primers from forward reads and quality control was performed using DADA2 (q2-dada2 plugin) [40]. The sequences were truncated to 250 bp based on the base call quality score during this step. After pre-processing steps, the resultant dataset contained 5,042,292 reads and amplicon sequences variant (ASV) table with associated sequences were generated from DADA2. Taxonomy classification was performed using q2-feature-classifier [41] with extracted 515-806 SILVA database [42] based on ASV table and associated sequences which were well-developed for bacterial data processing through QIIME2 following published protocols [39-42]. Mitochondria and chloroplast sequences were removed from the dataset resulting in 18,564 ASVs.

### 2.4 Data analysis

Both fungal and bacterial data were rarefied and analyzed using Phyloseq packages in R version 3.5.1 [ref 43] and Rstudio version 1.1.463 [ref 44] for taxonomic composition, alpha diversity, beta diversity, and endemic species [45]. Differences in alpha diversity were evaluated and compared using ANOVA with the “aov” function and pairwise multiple comparison (Tukey test) with the “TukeyHSD” function in R. Beta diversity was compared using PERMANOVA with the “adonis” function in the “vegan” package in R [46]. Network analysis was implemented with the SpiecEasi package [47] and followed the pipeline procedure for cross domain analysis [48]. Circular fungal-bacterial networks plots were generated using the “circlize” package in R [49]. We also performed indicator species analysis (function “indval” in “labdsv” package) in R [50] to identify the significant species at p = 0.05 that are predicted to be part of the structured crust types and sites.

## 3. Results

### 3.1 Geography, soil depth, and crust type influence biocrust microbial species richness

Alpha diversity analysis of biocrust microbial communities identified geographical differences for bacterial richness (ANOVA, F(3,20) = 4.845, p = 0.011) but not for fungal richness (ANOVA, F(3,20) = 1.493, p = 0.25) (Figure 2A). At JTNP, bacterial species richness was significantly lower than at GMT and KELSO, but not significantly different from the values at CIMA (Figure 2B). When comparing by soil depth, biocrust surface samples had significantly lower species richness than subsurface soil samples, both for fungal and bacterial richness (fungal t-test, t(43.594) = 3.208, p = 0.0025) (bacterial t-test, t(31.587) = 9.84, p = 3.856e-11) (Figure 2C and 2D). Also, fungal and bacterial richness of biocrust surface samples was significantly different by crust type (fungal ANOVA, F(4,19) = 9.456, p = 0.00022, bacterial ANOVA, F(4,19) = 4.256, p = 0.013 (Figure 2E). Fungal species richness in GLC was significantly lower than in other crust types. In addition, bacterial alpha diversity in LAC was significantly lower than in CLC, but not significantly lower than in other crust types (Figure 2F).

**Figure 2.**
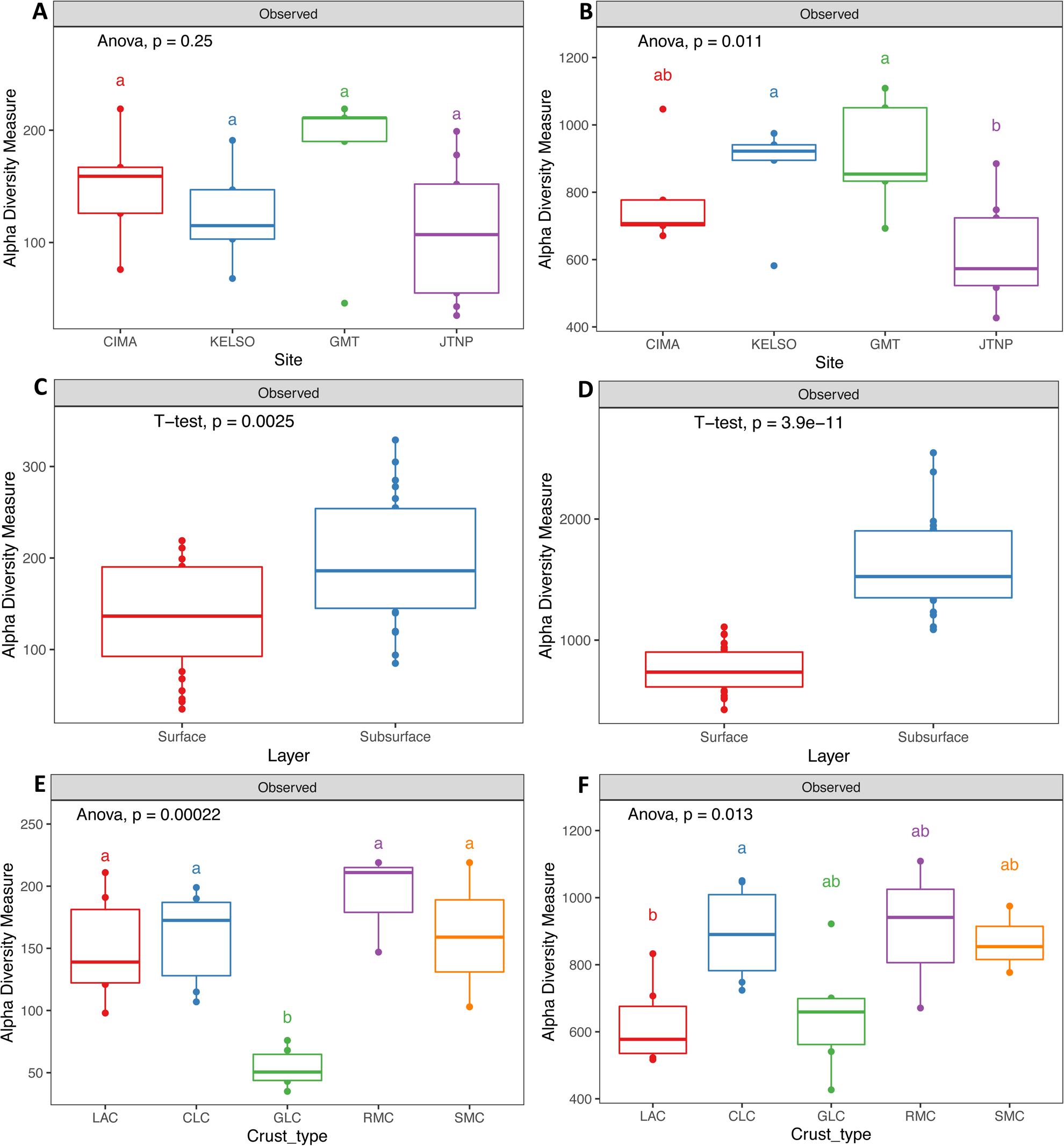
Boxplots showing alpha diversity as OTU richness in different site, soil depth, and crust type. A) Mojave biocrust fungal alpha diversity by site with rarefaction of 6842 reads per sample, B) Mojave bacterial alpha diversity by site with rarefaction of 37435 reads per samples, C) fungal alpha diversity by soil depth, D) bacterial alpha diversity by soil depth, E) fungal alpha diversity by crust types, and F) bacterial alpha diversity by crust type. Boxplots show 25th and 75th percentile while median was shown as lines inside boxes. Error bars show 1st and 99th percentile. Tukey HSD significant differences (P<0.05) are indicated by different letters.

There were 38 fungal taxonomic classes observed across all samples (Figure 3A and S1). We compared alpha diversity for each fungal class by site, layer, and crust type. Fungal richness comparison among sites showed significant differences within 3 fungal classes (Leotiomycetes (ANOVA, F(3,20) = 6.435, p = 0.00315), Blastocladiomycetes (ANOVA, F(3,20) = 4.818, p = 0.011), and Pucciniomycetes (ANOVA, F(3,20) = 3.912, p = 0.0239) (Figure S1). Generally, GMT had greater richness for these three fungal classes than the other sites while richness was lowest in JTNP. Blastocladiomycete and Pucciniomycete richness was greater in CIMA and GMT (central Mojave sites) than at JTNP, while Leotiomycetes richness was also greater at GMT and KELSO (central Mojave) than JTNP. When comparing by soil depth, 10 fungal classes showed significant differences richness (t-test, p < 0.05): Sordariomycetes, Agaricomycetes, Schizosaccharomycetes, Mucoromycetes, Saccharomycetes, Orbiliomycetes, Entomophthoromycetes, Mortierellomycetes, Basidiobolomycetes, and Pneumocystidomycetes (Table S1). All of these fungal classes were richer in subsurface soils than in the biocrust surface samples (Figure 3B, 3C, and 3D). Most fungal OTUs (514 OTUs) were shared between surface biocrust and subsurface samples (Figure S3B). Lastly, comparing fungal richness across crust types for each fungal class showed that 9 fungal classes were significantly different by crust type (ANOVA, p < 0.05): Sordariomycetes, Eurotiomycetes, Lecanoromycetes, Dothideomycetes, Leotiomycetes, Agaricomycetes, Schizosaccharomycetes, Pezizomycetes, and Tremellomycetes (Figure S1B and Table S2): GLC generally had lower fungal richness than the other crust types. Moss crusts (RMC and SMC) had greater fungal richness for Leotiomycetes, Peziozomycetes, and Tremellomycetes.

**Figure 3.**
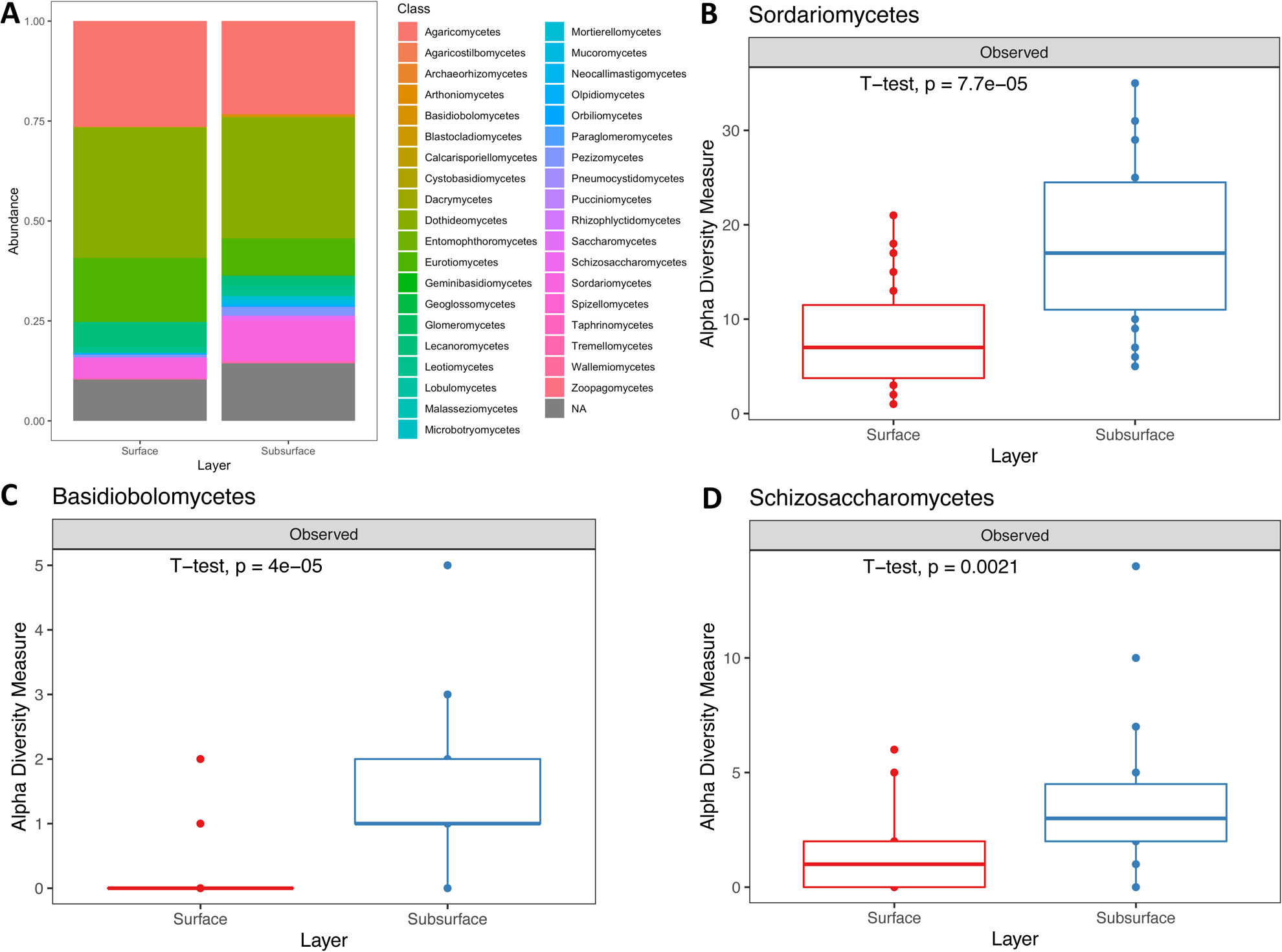
A) Fungal taxonomic composition bar plots at class level by layer (NA = unidentified), Top three fungal classes with significantly different alpha diversity by layer, including B) Sodariomycetes, C) Basidiobolomycetes, and D) Schizosaccharomycetes. Subsurface soil also had greater species richness than surface biocrust for seven other fungal taxonomic classes (Agaricomycetes, Mucoromycetes, Saccharomycetes, Orbiliomycetes, Entomophthoromycetes, Mortierellomycetes, and Pneumocystidomycetes). Boxplots show 25th and 75th percentile while median was shown as lines inside boxes. Error bars show 1st and 99th percentile. Tukey HSD significant differences (P<0.05) are indicated by different letters.

Bacterial alpha diversity was significantly different when comparing crust types, sites, and soil layers, and can be seen as differences in the relative abundance values in the taxonomic composition bar plot among the 30 prokaryotic phyla (Figure 4 and S2). Bacterial species richness comparison (for each phylum) by site indicated there was variable distribution of richness among 10 bacterial phyla (ANOVA, p < 0.05); including Proteobacteria, Firmicutes, Bacteroidetes, Actinobacteria, Acidobacteria, Planctomycetes, Patescibacteria, Armatimonadetes, Gemmatimonadetes, and Verrucomicrobia (Figure S2A and Table S3). In Proteobacteria, Firmicutes, Bacteroidetes, Actinobacteria, Acidobacteria, and Verrucomicrobia, bacterial species richness was lower at JTNP than at other sites. Acidobacteria, Planctomycetes, Patescibacteria, and Gemmatimonadetes had greater richness at GMT than at any other site. When comparing by soil depth, 19 phyla showed significant differences between surface vs. subsurface samples (t-test, p < 0.05); Proteobacteria, Firmicute, Actinobacteria, Euryarchaeota, Nanoarchaeota, Thaumarchaeota, Acidobacteria, Planctomycetes, Patescibacteria, Elusimicrobia, Armatimonadetes, Chloroflexi, Gemmatimonadetes, Entotheonellaeota, Cyanobacteria, Nitrospirae, FBP, Fibrobacteres, and Verrucomicrobia (Figure 4 and Table S4). Nearly all of these phyla showed greater species richness in subsurface soil than in biocrust samples. Cyanobacteria were the only bacterial phylum with significantly greater richness in biocrust surface samples than in subsurface soil. Majority of bacterial surface ASVs (2883 ASVs) were shared between surface biocrust and subsurface samples (Figure S3A). Ten phyla were significantly different by crust type (ANOVA, p < 0.05); Proteobacteria, Acidobacteria, Planctomycetes, Patescibacteria, Armatimonadetes, Deinococcus-Thermus, Chloroflexi, Cyanobacteria, FBP, and Verrucomicrobia (Figure S2B and Table S5). In Proteobacteria, Acidobacteria, Planctomycetes, Patescibacteria, and Verrucomicrobia, species richness was greater in moss crusts (RMC and SMC) than in other crust types. Richness of Armatimonadetes was lowest in LAC while Chloroflexi richness was highest in CLC. Lastly, Cyanobacteria richness was lower in moss crusts than in other crust types, versus highest in CLC.

**Figure 4.**
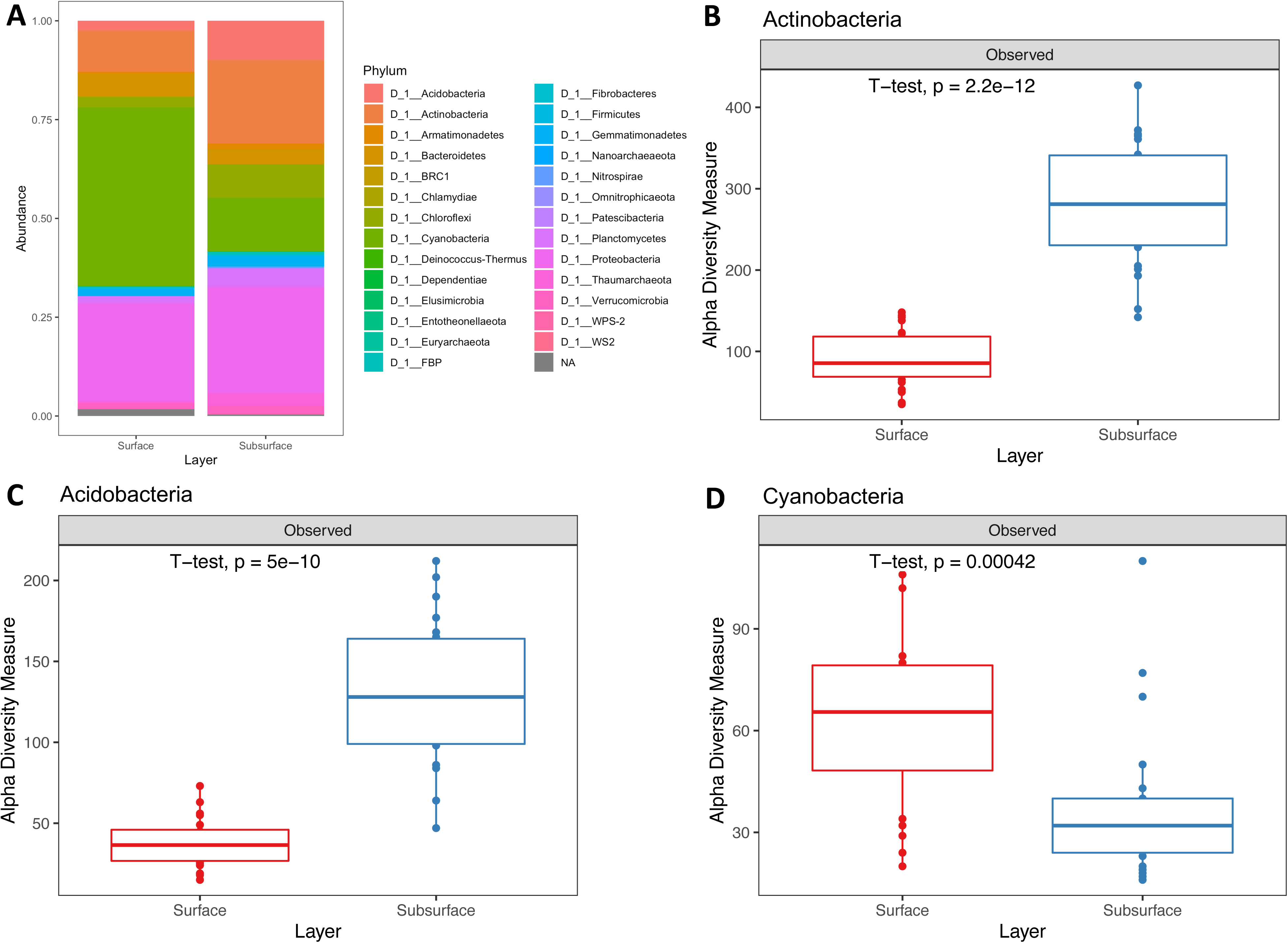
A) Bacterial taxonomic composition bar plot at phylum level by layer (NA = unidentified), Top two bacterial phyla in which alpha diversity by layer were significantly different including B) Actinobacteria and C) Acidobacteria. Same pattern was found in other 16 bacterial phyla in which subsurface soil had greater species richness than surface soil. D) Cyanobacteria bacterial richness on the soil surface was greater than in subsurface soil. Boxplots show 25th and 75th percentile while median was shown as lines inside boxes. Error bars show 1st and 99th percentile. Tukey HSD significant differences (P<0.05) are indicated by different letters.

### 3.2 Geography, soil depth, and crust type contribute to microbial composition pattern

Overall, the composition of biocrust fungal communities were significantly different when comparing by site (PERMANOVA, p = 0.001) and soil layer (PERMANOVA, p = 0.001) (Figure 5A). These differences in beta diversity were visualized in PCoA plots revealing 1) a geographical pattern: JTNP biocrust fungal composition clustered separately from central Mojave fungal communities (KELSO, GMT, and CIMA), 2) JTNP showed the strongest surface/subsurface clustering while central Mojave showed some surface/subsurface clustering, but not as clearly distinct as we observed in JTNP, and 3) Fungal beta diversity analysis showed that fungal community structures were also significantly different by crust type (PERMANOVA, p = 0.004) (Figure S4) but the clustering of samples by crust type was the weakest. GLC fungal community was distinctly different from other crust types, while the other biocrust types had overlaps in their fungal community composition,

**Figure 5.**
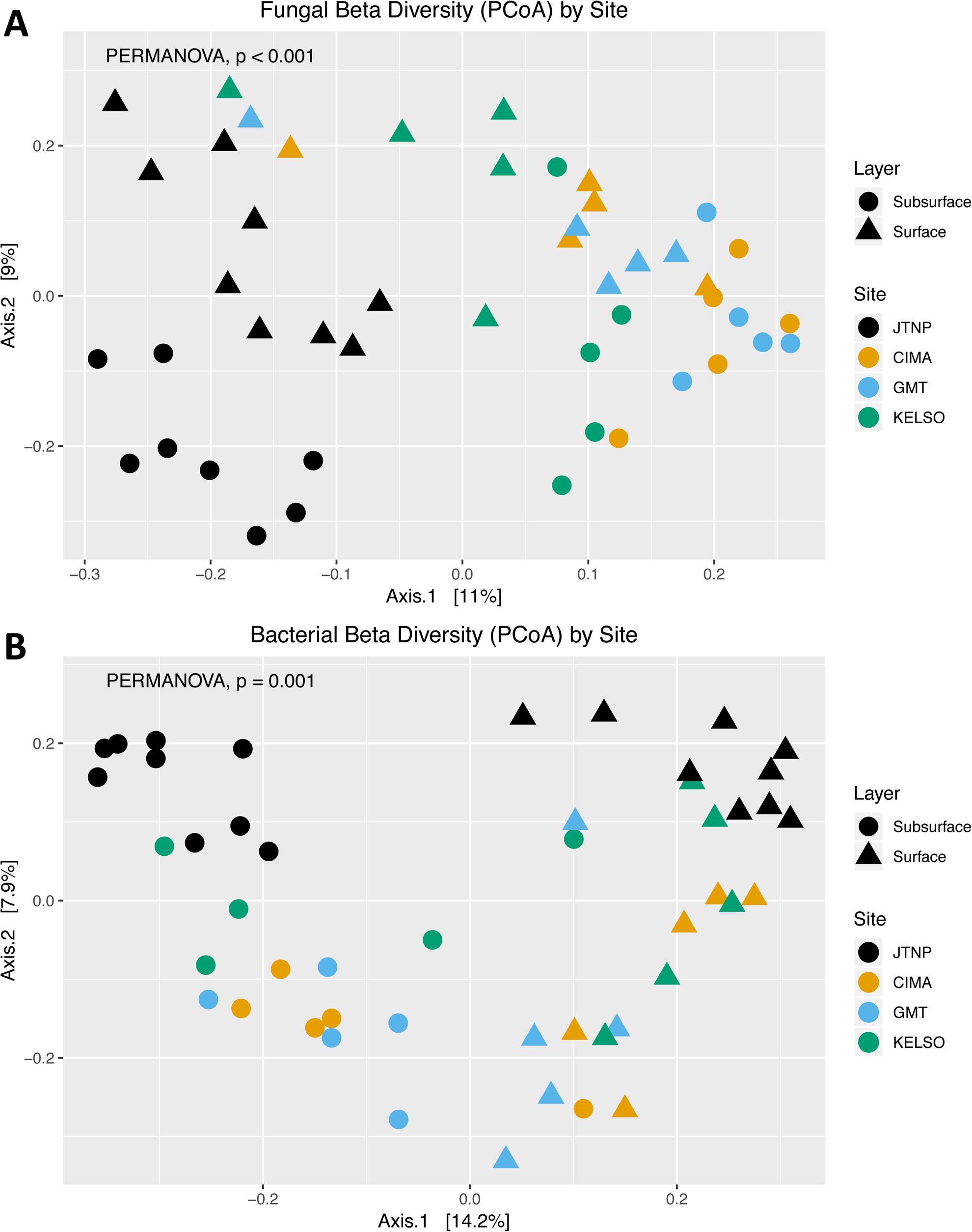
Beta diversity analysis of biocrust microbial communities. Dissimilarity of A) fungal and B) bacterial community composition in the comparison between site and soil depth (layer) using Principal Coordinate Analysis (PCoA). Different colors indicated four sampling sites including black color for JTNP, yellow color for CIMA, blue color for GMT, and green color for KELSO. Circle points showed subsurface samples while triangle points indicated surface samples. Significant differences (PERMANOVA, P < 0.05) were shown on PCoA plots.

Geography, soil depth, and biocrust types also influenced bacterial community composition. Bacterial communities were significantly different by site (PERMANOVA, p = 0.001) and layer (PERMANOVA, p = 0.001) (Figure 5B). Distinct clustering of JTNP bacterial communities away from the three sites of the central Mojave supported findings from the fungal communities. Bacterial communities also showed very strong surface/subsurface patterning where surface samples clustered closer together and the majority of subsurface samples were clustered near each other (nevertheless, two subsurface samples are clustered with surface samples). Moreover, when bacterial beta diversity was compared between different crust types, significant differences were detected (PERMANOVA, p = 0.001) (Figure S5) which produced a strong secondary pattern in addition to the surface/subsurface pattern (Figure 5B).

### 3.3 Fungal-bacterial networks underlie biocrust microbial connections

Microbial co-occurrence network analysis showed multiple discrete connections and abundance correlations among bacteria, archaea, and fungal biocrust communities. Bacterial networks were the most connected (∼54% were bacterial-bacterial connections) among all microbial communities both within and across domain networks (Figure 6). The network inferred from the abundances in surface samples indicated these communities are mostly structured within a large connected network (in the center of Figure 6) which contained Actinobacteria, Cyanobacteria, Proteobacteria, Ascomycota, and Basidiomycota as major backbone of microbial network hubs. For biocrust surface samples, cross domain fungal-bacteria links included; 1) Agaricomycetes and Dothideomycetes linked to Actinobacteria, 2) Agaricomycetes, Dothideomycetes, Eurotiomycetes, Orbilliomycetes, and Sordariomycetes linked to Cyanobacteria, 3) Dothideomycetes, Eurotiomycetes, and Pezizomycetes linked to Alphaproteobacteria, and 4) Dothideomycetes, Lecanoromycetes, and Sordariomycetes were linked to Blastocatellia (Figure 7). The complete network of microbial connections (both within and across domains) is depicted in Figure S6.

**Figure 6.**
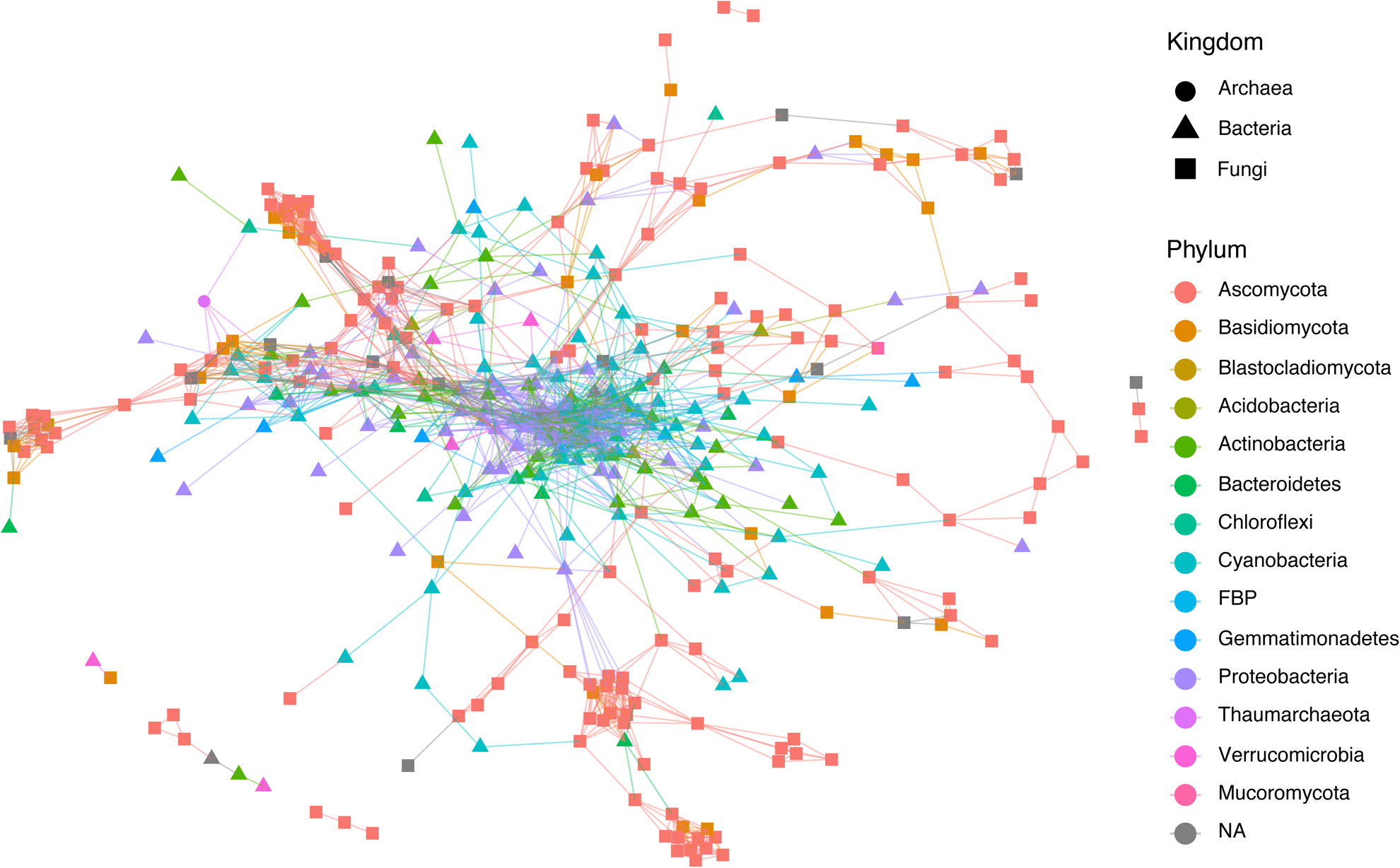
Microbial network analysis for biocrust surface samples. Each symbol/point on microbial network plot presents a single OTU. Microbial domains are indicated by different point shapes; archaea by circles, bacteria by triangles, and fungi by squares. Microbial networks are shown by line connection between points. Different colors indicate phylum for each point. Major network hubs concentrate at the center of microbial networks.

**Figure 7.**
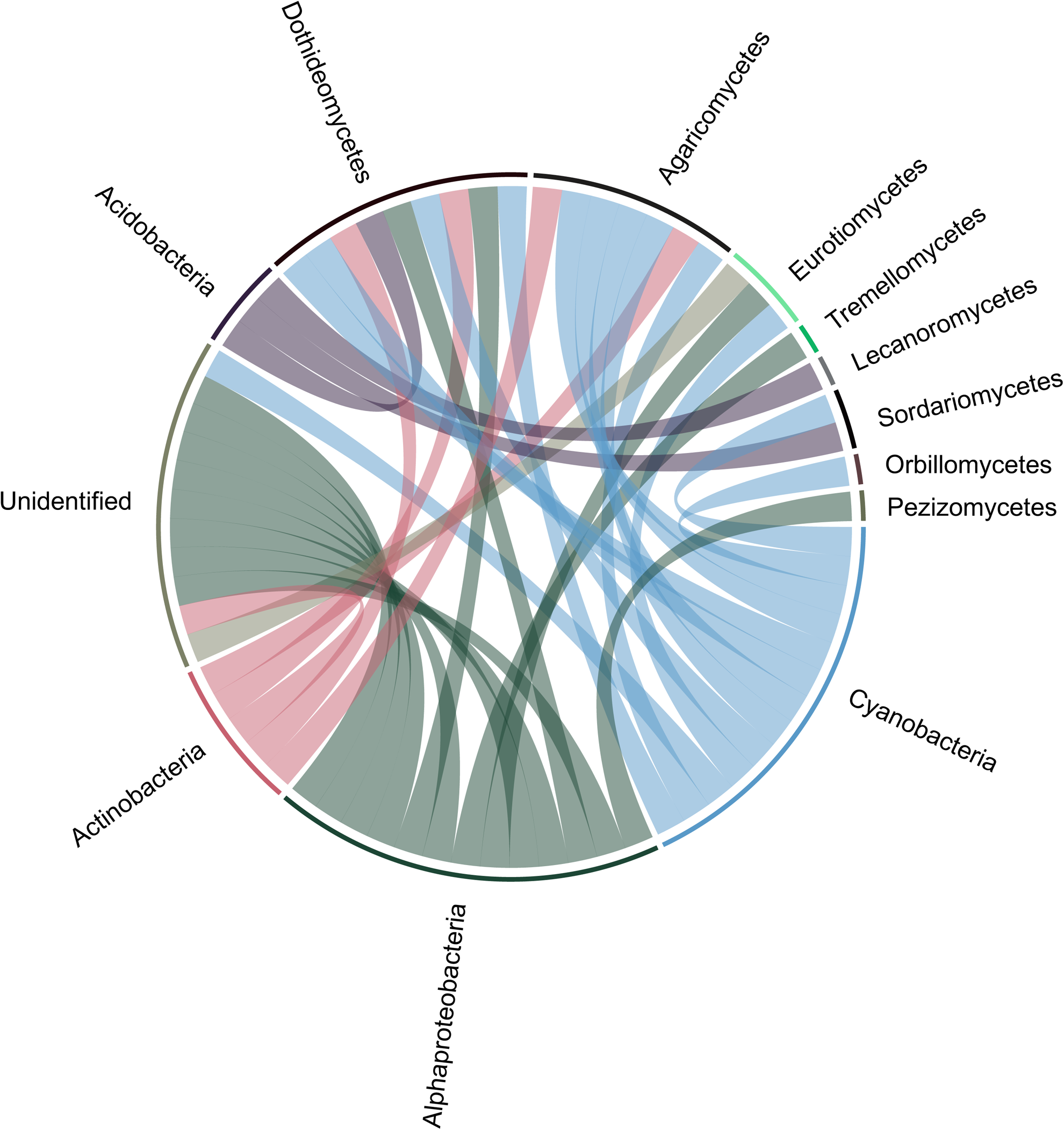
Cross-domain fungal-bacterial network analysis for biocrust surface samples. Cross-domain networks are a subset of total microbial networks showing in Figure 6. Each line represents the connection of a fungal OTU to a bacterial ASV. Different colors indicate different bacterial phyla. Cyanobacteria (in blue) had the highest number of connections to fungi. Cross-domain fungal-bacterial network analysis for biocrust subsurface samples are shown in Figure S7.

Subsurface soil samples showed similar patterns to surface biocrust where bacterial networks were more connected than other microbial community networks (Figure S7). Although large connected networks were observed as well, the major backbone and fungal-bacterial networks in subsurface communities revealed features different from surface microbial communities. The Mmajor phyla structuring subsurface microbial networks were Actinobacteria, Proteobacteria, Ascomycota, and Basidiomycota. Fungal-bacterial networks in subsurface samples included; 1) Agaricomycetes, Dothideomycetes, Eurotiomycetes, and Lecanoromycetes were linked to Actinobacteria, 2) Agaricomycetes, Basidiobolomycetes, Dothideomycetes, Leotiomycetes, Orbilliomycetes, and Sordariomycetes were linked to Alphaproteobacteria, 3) Dothideomycetes were linked to Bacteroidia, 4) Dothideomycetes and Mortierellomycetes were linked to Blastocatellia, 5) Dothideomycetes were linked to Chloroflexia, 6) Dothideomycetes were linked to Gammaproteobacteria, and 7) Eurotiomycetes were linked to Rubrobacteria (Figure S8).

### 3.4 Indicator and Endemic species specified geographical and crust type patterns

Overall, fungal indicator species analysis revealed fewer indicator species with lower abundance values than bacteria. Only 2 fungal indicator OTUs were detected by crust type and 11 species by site. Specifically, for RMC there was 1 species/OTU (closest related taxon: *Sporormia subticinensis*, dung saprotroph) and for SMC 1 species/OTU (closest related taxon: *Acrophialophora levis*, plant pathogen), but no fungal indicator species occurred in CLC, GLC, and LAC (Table S5). Fungal species by site revealed 2 indicator species/OTUs for CIMA (closest related taxon: *Catenulomyces convolutus*, unassigned functional guild and *Preussia terricola*, dung saprotroph and/or plant saprotroph), 4 indicator species/OTUs for JTNP (closest related taxon: *Allophoma labilis*, plant pathogen, *Curvularia inaequalis* plant pathogen, *Entoloma halophilum* (ectomycorrhizal, fungal parasite, and/or soil saprotroph), and *Preussia africana* (dung saprotroph and/or plant saprotroph), 5 indicator species/OTUs for KELSO (closest related taxon: *Alternaria hungarica* (animal pathogen, endophyte, plant pathogen, and/or wood saprotroph), *Cladosporium herbarum* (plant pathogen and/or wood saprotroph), *Colletotrichum gloeosporioides* (endophyte and/or plant pathogen), *Fusarium oxysporum* (plant pathogen, soil saprotroph, and/or wood saprotroph), and *Ulocladium dauci* (plant pathogen), but no fungal indicator species were observed at GMT (Table S7). In biocrust surface samples from all four sites, the top 50 endemic fungal species/OTUs (unique OTUs that were found specifically at each site) were mostly from Ascomycota, which included Acarosporales, Capnodiales, Chaetothyriales, Eurotiales, Geoglossales, Helotiales, Lecanorales, Orbilliales, Ostropales, Peltigerales, Pezizales, Pleosporales, Sordariales, Umbillicariales, and Verrucariales (Figure 8). Basidiomycota (Agaricales and Filobasidiales) and Mortierellomycota (Mortierellales) were also found to contain endemic species/OTUs. Identifiable fungal endemic OTUs were most similar to *Acarospora fuscata* (lichenized fungi: endemic to GMT), *Acrophialophora levis* (plant pathogen: endemic to KELSO), *Cladosporium herbarum* (plant pathogen and/or wood saprotroph: endemic to JTNP), *Curvularia affinis* (plant pathogen: endemic to JTNP), and *Deniquelata barringtoniae* (saprotroph: endemic to JTNP). Other endemic fungi were not identifiable at species level.

**Figure 8.**
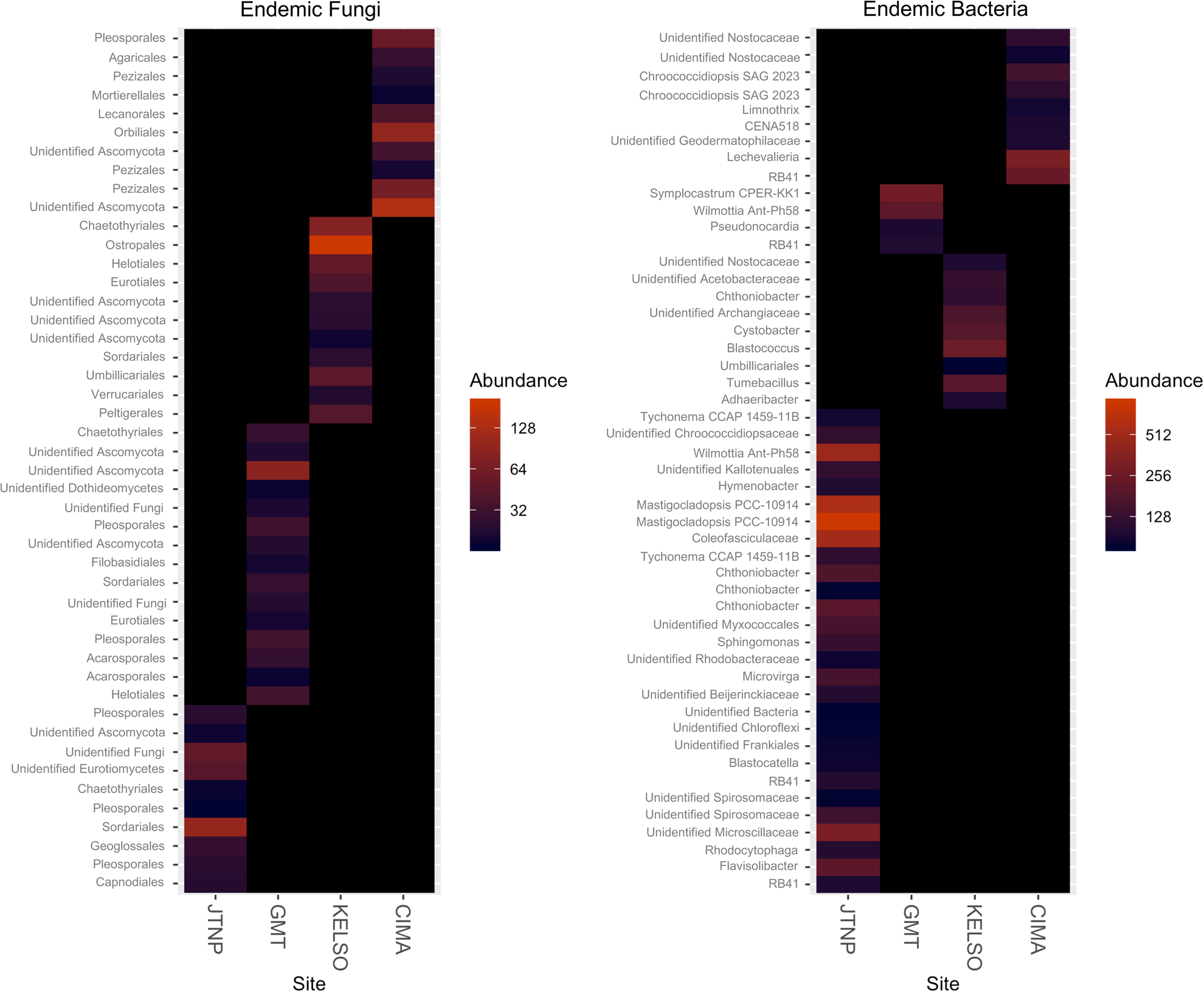
Heatmap for relative abundances of endemic fungal and bacterial species at each sampling site. (50 most abundant endemic OTUs/ASVs, identified as unique OTUs/ASVs found only at one site each). Each OTU/ASV is indicated by its most closely related named taxon on the left. Different color levels show relative abundances: red indicates high abundance and black indicates low abundance. Columns on the x-axis represent the four sampling sites.

In contrast, prokaryote indicator species analysis of biocrust surface samples showed 36 bacterial indicator ASVs within crust types and 67 ASVs when analyzed by site. There were 9 bacterial indicator species/ASVs for CLC (most closely similar to proteobacteria *Azospirillum soli*), 4 indicator species/ASVs for LAC (closest to proteobacteria *Belnapia moabensis*), 6 indicator species/ASVs for RMC (closest to proteobacteria *Salinarimonas sp. BN140002*) and 17 indicator species/ASVs for SMC (closest relatives were deinococcus-thermus *Deinococcus pimensis DSM 21231* and cyanobacteria *Calothrix sp. HA4186-MV5*), while no bacterial indicator species/ASVs were observed in GLC (Table S8). Bacterial indicator species/ASVs by site contained 29 indicator species/ASVs for CIMA (3 identifiable species/ASVs were most closely similar to bacteroidetes *Segetibacter aerophilus*, cyanobacteria *Chroococcidiopsis sp. BB79.2*, deinococcus-thermus *Deinococcus maricopensis DSM 21211*), 15 indicator species/ASVs for GMT (only 1 identifiable species, most closely similar to bacteroidetes *Parahymenobacter deserti*), 12 indicator species/ASVs for JTNP (1 identifiable species was bacteroidetes *Hymenobacter rigui*), and 11 indicator species/ASVs for KELSO (closest to proteobacteria *Roseomonas pecuniae*, proteobacteria *Sphingomonas kaistensis*) (Table S9). Bacterial endemic species showed a different pattern in which JTNP contained more bacterial endemics (28 species/ASVs) than all central Mojave sampling sites combined (22 species/ASVs). KELSO and CIMA had 9 bacterial endemics while GMT had the lowest number of bacterial endemics (4 species/ASVs). All endemic species/ASVs were from Acidobacteria, Actinobacteria, Bacteroidetes, Chloroflexi, Cyanobacteria, Firmicutes, Gemmatimonadetes, Proteobacteria, and Verrucomicrobia. Identifiable bacterial endemic genera included ASVs which were most closely similar to *Adhaeribacter* (bacteroidetes), *Blastocatella* (acidobacteria), *Blastococcus* (actinobacteria), *Chroococcidiopsis* (cyanobacteria), *Chthoniobacter* (verrucomicrobia), *Cystobacter* (proteobacteria), *Flavisolibacter* (bacteroidetes), *Gemmatirosa* (gemmatimonadetes), *Hymenobacter* (bacteroidetes), *Lechevalieria* (actinobacteria), *Limnothrix* (cyanobacteria), *Mastigocladopsis* (cyanobacteria), *Microvirga* (proteobacteria), *Pseudonocardia* (actinobacteria), *Rhodocytophaga* (bacteroidetes), *Sphingomonas* (proteobacteria), *Symplocastrum* (cyanobacteria), *Tumebacillus* (firmicutes), *Tychonema* (cyanobacteria), and *Wilmottia* (cyanobacteria) (Figure 8).

## 4. Discussion

In our study, we detected several distinct patterns structuring biocrust microbial communities in the Mojave Desert. These patterns include 1) a strong geographical differentiation between the three central Mojave sites (GMT, KELSO, and CIMA) versus the southern Mojave site (JTNP), 2) a soil depth pattern which clearly differentiates biocrust surface diversity from subsurface microbial communities, and 3) a biocrust type pattern which shows differences between LAC, CLC, GLC, SMC, and RMC. Microbial network analysis showed Cyanobacteria as the major hubs for overall biocrust microbial community connectivity, while Pleosporales were the major fungal hubs for fungal-bacteria networks. Recognizing these patterns, continuing to disentangle their biotic linkages, and investigating how these patterns and networks affect ecosystem functions will contribute crucial information to dryland conservation and restoration management.

### 4.1 *Geographical pattern:* Does geographical location structure biocrust microbial communities and reveal unique microbial species?

Previous research has identified characteristics that determine biocrust distribution both globally and locally which included “biogeographic, climatic, edaphic, topographic, and biotic” factors [51]. Geographical distribution of these biocrusts was previously examined using externally visible morphology for scoring crust coverage fractions and occurrence across the region of study. Many previous case studies have demonstrated biogeography based on presence/absence of functional biocrust types, but these can be difficult to compare across several studies due to differences in sampling methods and taxonomic keys used, or by focusing on single biocrust type [51,52]. In our study, we employed both morphological identification and whole community microbial NGS to determine the regional distribution of biocrust. We observed distinct geographical patterns within the Mojave Desert in which both alpha and beta diversity differentiated microbial communities in central Mojave (GMT, KELSO, and CIMA) from southern Mojave (JTNP). Our results are similar to previous studies in which microbial communities were more similar when collecting sites were in close proximity [30-32]. However, our findings also indicate that major differences were likely due to a Mojave-Colorado desert ecotone effect at JTNP, which significantly separated southern Mojave microbial communities from central Mojave diversity.

Despite us sampling the same functional biocrust types with similar external morphologies in all 4 localities, beta diversity and indicator species analysis indicated that central Mojave Desert localities had unique microbial communities in the surveyed crust types dissimilar from the same crust types sampled in JTNP. Although external morphology was not visibly different, our results demonstrated that microbial communities and indicator microorganisms showed greater resolution to identify geographical pattern/separation at the Mojave-Colorado desert ecotone than morphological identification. Similar procedures previously identified a boundary of Mojave desert and Colorado desert using vascular plant taxon composition and key species [53]. Employing a similar procedure, we identified major fungal and bacterial components that contributed significantly to biocrust microbial geographical patterning (ANOVA, p < 0.05). These included two phyla in fungal communities (Sordariomycetes and Agaricomycetes) and eleven phyla in bacterial communities (Proteobacteria, Bacteroidetes, Actinobacteria, Acidobacteria, Planctomycetes, Armatimonadetes, Chloroflexi, Gemmatimonadetes, Cyanobacteria, FBP, and Verrucomicrobia). To our knowledge, this is the first report of biocrust geographical fungal indicator species. They are plant associated fungi specifically found in biocrusts at each site, for example: *Allophoma labilis* -plant pathogen (JTNP) [25,54]; *Curvularia inaequalis* -Plant pathogen (JTNP) [25,55]; *Cladosporium herbarum -*Plant Pathogen and/or Wood Saprotroph (KELSO) [25,56,57]; *Colletotrichum gloeosporioides -*Endophyte and/or Plant Pathogen (KELSO) [25,56]; *Ulocladium dauci -*Plant Pathogen (KELSO) [25,56]; and *Preussia terricola -*Dung Saprotroph and/or Plant Saprotroph (CIMA) [25,58]. Although the fungal loop hypothesis is a major focus for plant-biocrust interaction studies [59], our results suggest other important plant-biocrust interactions are mediated through the fungal community. Biocrust ability to capture seeds [60] could also mean that biocrusts are able to trap plant associated fungal spores, which then develop roles within the biocrust community for decomposition purposes.

### 4.2 *Soil depth pattern:* Do biocrust microbial surface communities differ from those in the underlying soil subsurface?

As a living skin of drylands, biocrusts are usually studied with exclusive focus on the surface [2]. As an example of the few exceptions, pioneering biocrust work on soil depth patterns was conducted on the Colorado Plateau, using culture-based quantification of viable aerobic copiotrophs (VAC) to show that bacterial populations in biocrust compared to its own subsurface soil or to soil without crust [8]. However, DGGE fingerprints of bacterial community in the same study did not show soil depth as a factor in microbial diversity patterning. Bacterial communities were subsequently shown soil depth pattern in Colorado Plateau and central Mojave using NGS approach which provided more resolution to microbial diversity [28,32]. Only one study to date has surveyed both the fungal and bacterial communities in and below biocrusts, reporting soil depth patterning in southern Nevada with higher subsurface diversity for both domains [33]. In general, our soil depth pattern results were similar to previous studies when we explored high-level taxonomic diversity and species richness. We found lower overall species richness within biocrust samples than in subsurface soil. Cyanobacteria on the other hand were greater in species richness for biocrust than in subsurface soil, due to their dependence on availability of light [8, 28]. In previous studies, the major bacterial phyla (Acidobacteria, Actinobacteria, Chloroflexi, and Proteobacteria) were observed to be more diverse and had greater richness in subsurface soil [28,32]. This confirmed our results, and we furthermore found that 12 additional bacterial and 3 archaeal phyla were also significantly different between surface vs. subsurface soil in our data set (Table S4). Thus, Mojave biocrust bacterial communities showed more vertical differences than previous publications, which may indicate greater differentiation of functional traits. We also explored the fungal community, which indicated similar pattern to the bacterial community. Fungal species richness in biocrust surface samples was greater than in subsurface soil. We also found that Sordariomycetes had significant soil depth pattern similar to a previous study in Nevada [33]. However, nine additional fungal classes showed significant soil depth pattern at our collection sites in Mojave desert, California compared to southern Nevada [33].

To our knowledge, this study is the first biocrust microbial study, which incorporated cross domain (fungal-bacterial) networks. Incorporating fungal with bacterial community in microbial network analysis improves network stability compared to single-domain microbial networks [48]. To better understand the entire microbial network in biocrust systems, bacteria and fungi were jointly analyzed in a single cross-domains network analysis, identifying key microorganisms in both domains. Our network analysis showed that Cyanobacteria were the key microbial hubs for biocrusts (surface), which corresponds to previous findings [6,16] and to their high biomass previously detected in biocrust surface samples [8,28,33]. We also identified Pleosporales (Dothideomycetes) as the key to fungal-bacterial connections, which has been previously shown as dominant fungal taxa in biocrust and semiarid area [31,61]. We found that Agaricomycetes were another major fungal hub, which fits the abundance of the group as reported in the southern Nevada study [33]. Thus, our data suggested two groups of fungi that could potentially be key microorganisms for biocrusts in both southern California and Nevada. While Cyanobacteria were not present as a hub in our analyses of subsurface soil samples, Pleosporales (Dothideomycetes) were also a major hub for fungal-bacterial networks in subsurface soil while Agaricomycetes were also found as a minor hub. Therefore, our results indicated that fungal hubs were similar between surface and subsurface, while bacterial networks were different mostly because of photoautotrophs. Overall, we noted a strong soil depth pattern in our Mojave biocrusts, with greater numbers of bacterial phyla and fungal classes contributing to these patterns than previously reported [8,28,33]. Nonetheless, functional guild could not be identified for the fungi that contributed to soil depth patterning. If functional guild was identifiable, it could potentially provide essential information of the fungal hubs.

### 4.3 *Biocrust type pattern:* Are biocrust types linked with microbial diversity?

Different biocrust types have been well studied and are classified by a combination of their morphology, aggregation strength and by dominant photoautotrophic community members such as cyanobacterial or algal, bryophytes, and lichens [2,3,19,20]. Many environmental factors such as temperature, moisture, salinity, soil texture, dust deposition, etc. influence the different crust types that occur in the region [2]. However, we have very little knowledge about how biocrust types are linked with their microbial communities. In order to fill in this gap, we examined and compared five different biocrust types including LAC, CLC, GLC, SMC, and RMC for both bacterial and fungal community. We observed lower bacterial species richness in LAC, which was expected because LAC is an initial phase of biocrust formation. These results were similar to previous findings, which indicated lower species richness in early stage biocrust [3,62]. However, these differences were not observed in fungal communities, which suggests very small changes in fungal richness occur through biocrust successional processes. Fungal alpha diversity showed similar patterns across all biocrust types, except for GLC which had the lowest fungal species richness, indicating that some fungi were relatively more abundant in this crust type (Figure S1B). Therefore, we have shown that fungal and bacterial communities did not necessarily follow the same trend in structuring their communities in different crust types.

Cyanobacteria, which was previously indicated to be the major microbial hubs from our network analysis, showed that they were more abundant in LAC, CLC, and GLC than in SMC and RMC. The presence of Cyanobacteria in LAC and lichen crusts, but not in moss crusts is indicative of their central role as primary autotrophic community members. Alpha diversity analysis also differentiated LAC, CLC, GLC from moss crusts (SMC and RMC) in their composition of Proteobacteria, Actinobacteria, and Acidobacteria. In these phyla, greater alpha diversity was observed in moss crusts than LAC and lichen crusts. Moss crust has been shown to retain more moisture than light cyanobacterial crust [63] and fixes carbon at higher rates [4], thus contributing greater quantities of organic matter to support other soil bacteria. In addition, our indicator species analysis also showed that both types of moss crust contained both fungal and bacterial indicator species (Table S6 and S8) indicating that moss crusts could be distinguished by indicator species (we noted that indicator species may exist for each crust type but we recommend a higher sampling effort to test for this.). However, we were not able to identify identical lichen species from sequences as we described from external morphology, but several OTUs matched Peltigerales (with high abundance in CLC) which could possibly be *Collema sp.* and many OTUs matched Verrucariales (with high abundance in GLC) that might be *Clavascidium sp*. as we identified from morphology. This issue clearly shows that better molecular markers are needed for these lichens. Although a clear crust type pattern was not observed from indicator species analysis, we showed that crust type differences were detected with indicator species. Therefore, our results supported the crust types differences hypothesis that microbial communities are structured by crust types.

### 4.4 Implication to conservation and restoration management

Current efforts are attempting to restore biocrusts in heavily disturbed landscapes [64-67] are often met with little or no success. Our findings provide several implications for conservation and restoration management that have yet to be incorporated. In our dataset, which was limited to five biocrust types from four sampled sites, microbial communities from the same biocrust type in different locations are not identical and should therefore not be considered as interchangeable biocrust inoculum for restoration. Further, the efforts of establishing biocrust restoration from inoculum often focus on promoting biomass growth of primarily photoautotrophs, but much less on other biocrust microorganisms which could also be important biocrust components. Our data support that other biocrust microbes could also be essential components to these communities and we have shown that complex linkages within and between the two surveyed microbial domains, hub taxa, and indicator species do occur and are discoverable by the applied methods. However, although we discovered fungal hub taxa in our data set, the functional guild and community function of these species are still unclear. As a result, more research is still needed on the functional roles of desert soil fungi and how they may affect biocrust microbial communities before inoculation experiments can be designed. More sites and crust types within the Mojave Desert need to be studied and efforts to implement restoration methods like inoculation should be preceded by combined domain NGS surveys like the present work. This is especially true in drylands for which no baseline work on microbial diversity has been completed yet.

Considering our findings, we recommend the following steps when considering restoring biocrust with a goal of accounting for a more comprehensive microbial composition. In cases where conservation action is needed but monetary support is lacking, passive rehabilitation remains the slowest but less invasive strategy as it is the least likely to disrupt established source biocrust communities, allows for natural recovery, and thus most likely to yield sustainable outcomes [68,69]. In areas with urgent restoration needs, the ecoregion first must be sampled across its geographical extent to understand biocrust microbial diversity in the area and to explore factors which might contribute to biogeographical distribution. For example, previous research has shown biogeography of biocrusts and distribution of some key species was explainable in part by differences in temperature regimes [30-32,70]. Therefore, management programs should consider how biocrust restoration might affect these local microbial communities geographically. Especially, the risk of potentially distributing likely pathogens in the area may need closer investigation, as our results indicated some local fungal plant pathogens are presented in Mojave biocrusts which could possibly be dispersed by restoration procedure used. Second, certain microorganisms could be key members of the communities representing major hubs for microbial networks that have not been identified yet. Some of these microorganisms can be found in the soil beneath the crust or are shared between subsurface soil and biocrusts, suggesting that local subsurface soil should also be considered as starting material for biocrust production or inoculation. This strategy is consistent with a previous study that showed small shifts (in contrast to significantly change cyanobacteria composition) in the cyanobacterial community when using local soil/biocrust inoculum [67]. Lastly, we have also shown that biocrust microbial communities are different depending on the crust’s primary photoautotrophs (moss vs. lichen crusts) and external morphology should therefore be clearly identified. Likewise, other major knowledge gaps still exist in our understanding of the temporal variability of biocrust communities. For instance, we need a better understanding of how temporal changes and seasonality affect the microbes of resident biocrust communities, and how this may render those communities more or less suited for active microbial inoculation.

## 5. Conclusion

In summary, our findings provide the most extensive characterization of local biocrust microbiota to date from the central and southern Mojave Desert. It is to our knowledge the first comprehensive biocrust microbial community investigation, which revealed geographical, soil depth, and crust type diversity patterns considering both fungi and bacteria microbes. Although identification of biocrust types by their external morphology is practical for preliminary observation in the field, we have shown that microbial components within each type can be distinct geographically. Biocrust surface and subsurface communities also have unique microbial compositions. Our results supported the hypothesis that Cyanobacteria are key microorganisms in the biocrust with network analysis demonstrating that they are the major hubs for overall biocrust microbial community connectivity. We also identified Pleosporales as a major hub for fungal-bacteria networks. Our key findings imply that microbial species composition need to be taken into account in future biocrust restoration and management which should consider using either a passive restoration strategy or finding local inoculum to maintain and restore local biocrust microbial diversity. At a potential restoration site, initial assessment should also aim to understand spatial variation in the site’s microbial composition, including identification of potential keystone species and surveying for potential endemics. Neglecting these differences could possibly lead to unknown consequences to both biocrust microbial communities and desert ecosystems such as risk of pathogen spread, potential loss of microbial diversity and endemic species, and possibly destruction of biocrusts.

## Acknowledgements

We thank Joshua Tree National Park (Permit JOTR-2017-SCI-0010), the Mojave National Preserve (Permit MOJA-2015-SCI-0017), the University of California Natural Reserve System, and the Sweeney Granite Mountains Desert Research Center (SGNDRC) for permission to conduct research (DOI: 10.21973/N3S942). We thank Aurapat Ngamnithiporn, Derreck Carter-House, and Sangsan Warakkagun for assistance with biocrust sampling and transportation. We also gratefully acknowledge logistical support and central Mojave site selection guidance from UC reserve managers Tasha La Doux and Jim André at the SGNDRC, as well as National Parks archeologist David Nichols at MNP. Funding was provided by the USDA Agriculture Experimental Station at the University of California, Riverside and NIFA Hatch project CA-R-PPA-5062-H to J.E.S.; the California Desert Research Fund at The Community Foundation awarded to N. Pietrasiak; Robert Lee Graduate Student Research Grants awarded to N. Pombubpa and N. Pietrasiak; and a Mycological Society of America (MSA) Translational Mycology award to N. Pombubpa. Fungal ITS and bacterial 16S primer sequences and arrayed barcodes were provided by the Alfred P. Sloan Foundation Indoor Microbiome Project. Data analyses were performed on the High-Performance Computing Cluster at the University of California-Riverside in the Institute of Integrative Genome Biology supported by NSF DBI-1429826 and NIH S10-OD016290. N. Pombubpa was supported by the Royal Thai government fellowship.

## Supplemental Figure Legends

**Figure S1.** A) Fungal taxonomic composition bar plots at class level by sites, Top three fungal classes with significantly different alpha diversity by sites, including Leotiomycetes, Blastocladiomycetes, and Pucciniomycetes.B) Fungal taxonomic composition bar plot at class level by crust types, top three fungal classes in which alpha diversity by layer was significantly different including Dothideomycetes, Eurotiomycetes, and Sordariomycetes. Boxplots show 25th and 75th percentile while median was shown as lines inside boxes. Error bars show 1st and 99th percentile.

**Figure S2.** A) Bacterial taxonomic composition bar plot at phylum level by sites, three bacterial phyla in which alpha diversity by layer was significantly different including Acidobacteria, Actinobacteria, and Verrucomicrobia (ANOVA, p < 0.05). B) Bacterial taxonomic composition bar plot at phylum level by crust types, top three bacterial phyla in which alpha diversity by layer was significantly different including Cyanobacteria, Proteobacteria, and Acidobacteria (ANOVA, p < 0.05).

**Figure S3.** A) Bacterial surface vs. subsurface; sharing 2883 ASVs between surface biocrust and subsurface samples. B) Fungal surface vs. subsurface; sharing 514 OTUs between surface biocrust and subsurface samples.

**Figure S4.** Fungal Beta Diversity (PCoA) by Crust type, showing that GLC fungal composition was significantly different thfroman other crust types collected in all 3 Mojave desert sites (CLC, LAC, RMC, and SMC) (PERMANOVA, p = 0.004).

**Figure S5.** Bacterial Beta Diversity (PCoA) by Crust type showing that crust types differed significantly in the degree of divergence between surface versus subsurface community in the ordination plots (PERMANOVA, p = 0.001).

**Figure S6.** Microbial networks analysis for biocrust surface samples showed that bacterial networks were more connected than archaeal networks, fungal networks and fungal-bacterial networks.

**Figure S7.** Microbial network analysis for biocrust subsurface samples showed that bacterial networks were more connected than archaea networks, fungal networks and fungal-bacterial networks.

**Figure S8.** Cross-domain fungal-bacterial networks analysis for subsurface samples showed that Actinobacteria, Proteobacteria (Alphaproteobacteria) and Ascomycota (Dothideomycetes) were the major hubs for subsurface microbial networks.

## References

1. Belnap J, Büdel B, Lange OL (2001) Biological soil crusts: characteristics and distribution. In: Biological soil crusts: structure, function, and management. Springer, Berlin, Heidelberg, pp 3–30

2. Belnap J, Weber B, Büdel B (2016) Biological soil crusts as an organizing principle in drylands. In: Biological soil crusts: an organizing principle in drylands. Springer, Cham, pp 3–13

3. Maier S, Tamm A, Wu D, Caesar J, Grube M, Weber B (2018) Photoautotrophic organisms control microbial abundance, diversity, and physiology in different types of biological soil crusts. The ISME journal 12:1032. https://doi.org/10.1038/s41396-018-0062-8

4. Pietrasiak N, Regus JU, Johansen JR, Lam D, Sachs JL, Santiago LS (2013). Biological soil crust community types differ in key ecological functions. Soil Biology and Biochemistry 65:168–171. https://doi.org/10.1016/j.soilbio.2013.05.011

5. Weber B, Belnap J, Büdel B (2016) Synthesis on Biological Soil Crust Research. In: Biological Soil Crusts: An Organizing Principle in Drylands. Springer International Publishing, pp 527–534

6. Belnap J, Gardner JS (1993) Soil microstructure in soils of the Colorado Plateau: the role of the cyanobacterium Microcoleus vaginatus. Great Basin Naturalist 53:6.

7. Garcia-Pichel F, López-Cortés A, Nübel U (2001) Phylogenetic and morphological diversity of cyanobacteria in soil desert crusts from the Colorado Plateau. Appl. Environ. Microbiol. 67:1902–1910. https://doi.org/10.1128/AEM.67.4.1902-1910.2001

8. Garcia-Pichel F, Johnson SL, Youngkin D, Belnap J (2003) Small-scale vertical distribution of bacterial biomass and diversity in biological soil crusts from arid lands in the Colorado Plateau. Microbial Ecology 46:312–321. https://doi.org/10.1007/s00248-003-1004-0

9. Gundlapally SR, Garcia-Pichel F (2006) The community and phylogenetic diversity of biological soil crusts in the Colorado Plateau studied by molecular fingerprinting and intensive cultivation. Microbial Ecology 52:345–357. https://doi.org/10.1007/s00248-006-9011-6

10. Maier S, Muggia L, Kuske CR, Grube M (2016) Bacteria and Non-lichenized Fungi Within Biological Soil Crusts. In: Biological Soil Crusts: An Organizing Principle in Drylands. Springer International Publishing, pp 81–100

11. Fritsch FE, John RP (1942) An Ecological and Taxonomic Study of the Algae of British Soils: II. Consideration of the Species observed. Annals of Botany 6:371–395.

12. Hallmann C, Stannek L, Fritzlar D, Hause-Reitner D, Friedl T, Hoppert M (2013) Molecular diversity of phototrophic biofilms on building stone. FEMS Microbiology Ecology 84:355–372. https://doi.org/10.1111/1574-6941.12065

13. Hoppert M, Reimer R, Kemmling A, Schröder A, Günzl B, Heinken T (2004) Structure and reactivity of a biological soil crust from a xeric sandy soil in Central Europe. Geomicrobiology Journal 21:183–191. https://doi.org/10.1080/01490450490275433

14. Komarek J (2013) Cyanoprokaryota: Heterocytous Genera. Springer Spektrum. 3rd Part. In: Büdel B, Gärtner G, Krienitz L, Schagerl M Süßwasserflora von Mitteleuropa, Bd. 19 (3), Springer Spektrum, Berlin, Heidelberg, pp 1–1130

15. Ruprecht U, Brunauer G, Türk R (2014) High photobiont diversity in the common European soil crust lichen Psora decipiens. Biodiversity and conservation 23:1771–1785. https://doi.org/10.1007/s10531-014-0662-1

16. Büdel B, Dulić T, Darienko T, Rybalka N, Friedl T (2016) Cyanobacteria and algae of biological soil crusts. In: Biological soil crusts: an organizing principle in drylands. Springer, Cham, pp 55–80

17. Amann RI, Ludwig W, Schleifer KH (1995) Phylogenetic identification and in situ detection of individual microbial cells without cultivation. Microbiol. Mol. Biol. Rev. 59:143–169.

18. Viaud M, Pasquier A, Brygoo Y (2000) Diversity of soil fungi studied by PCR–RFLP of ITS. Mycological Research 104:1027–1032. https://doi.org/10.1017/S0953756200002835

19. Bowker MA, Belnap J, Miller ME (2006) Spatial modeling of biological soil crusts to support rangeland assessment and monitoring. Rangeland Ecology & Management 59:519–529. https://doi.org/10.2111/05-179R1.1

20. Büdel B, Darienko T, Deutschewitz K, Dojani S, Friedl T, Mohr KI, Salisch M, Reisser W and Weber B (2009) Southern African biological soil crusts are ubiquitous and highly diverse in drylands, being restricted by rainfall frequency. Microbial ecology 57:229–247. https://doi.org/10.1007/s00248-008-9449-9

21. Bates ST, Garcia□Pichel F (2009) A culture□independent study of free□living fungi in biological soil crusts of the Colorado Plateau: their diversity and relative contribution to microbial biomass. Environmental Microbiology 11:56–67. https://doi.org/10.1111/j.1462-2920.2008.01738.x

22. Steven B, Yeager C, Belnap J, Kuske CR (2014) Common and distinguishing features of the bacterial and fungal communities in biological soil crusts and shrub root zone soils. Soil Biology and Biochemistry 69:302–312. https://doi.org/10.1016/j.soilbio.2013.11.008

23. Steven B, Hesse C, Gallegos-Graves LV, Belnap J, Kuske CR (2015) Fungal diversity in biological soil crusts of the Colorado plateau. In: Proc 12th Biennial Conf Science Management Colorado Plateau

24. Smith DP, Peay KG (2014) Sequence depth, not PCR replication, improves ecological inference from next generation DNA sequencing. PLoS One 9:90234. https://doi.org/10.1371/journal.pone.0090234

25. Nguyen NH, Song Z, Bates ST, Branco S, Tedersoo L, Menke J, Schilling JS, Kennedy PG (2016) FUNGuild: an open annotation tool for parsing fungal community datasets by ecological guild. Fungal Ecology 20:241–248. https://doi.org/10.1016/j.funeco.2015.06.006

26. Peay KG, Kennedy PG, Talbot JM (2016) Dimensions of biodiversity in the Earth mycobiome. Nature Reviews Microbiology 14:434. https://doi.org/10.1038/nrmicro.2016.59

27. Caporaso JG, Lauber CL, Walters WA, Berg-Lyons D, Lozupone CA, Turnbaugh PJ, Fierer N, Knight R (2011) Global patterns of 16S rRNA diversity at a depth of millions of sequences per sample. Proceedings of the National Academy of Sciences 108:4516–4522. https://doi.org/10.1073/pnas.1000080107

28. Steven B, Gallegos-Graves LV, Belnap J, Kuske CR (2013) Dryland soil microbial communities display spatial biogeographic patterns associated with soil depth and soil parent material. FEMS Microbiology Ecology 86:101–113. https://doi.org/10.1111/1574-6941.12143

29. Liu L, Liu Y, Zhang P, Song G, Hui R, Wang Z, Wang J (2017) Development of bacterial communities in biological soil crusts along a revegetation chronosequence in the Tengger Desert, northwest China. Biogeosciences 14:3801–3814. https://doi.org/10.5194/bg-14-3801-2017

30. Nagy ML, Pérez A, Garcia-Pichel F (2005) The prokaryotic diversity of biological soil crusts in the Sonoran Desert (Organ Pipe Cactus National Monument, AZ). FEMS Microbiology Ecology 54:, 233-245. https://doi.org/10.1016/j.femsec.2005.03.011

31. Bates ST, Nash III TH, Garcia-Pichel F (2012) Patterns of diversity for fungal assemblages of biological soil crusts from the southwestern United States. Mycologia 104:353–361. https://doi.org/10.3852/11-232

32. Mogul R et al. (2017) Microbial community and biochemical dynamics of biological soil crusts across a gradient of surface coverage in the Central Mojave Desert. Frontiers in microbiology 8:1974. https://doi.org/10.3389/fmicb.2017.01974

33. Mueller RC, Belnap J, Kuske CR (2015) Soil bacterial and fungal community responses to nitrogen addition across soil depth and microhabitat in an arid shrubland. Frontiers in microbiology, 6:891. https://doi.org/10.3389/fmicb.2015.00891

34. Palmer JM, Jusino MA, Banik MT, Lindner DL, (2018) Non-biological synthetic spike-in controls and the AMPtk software pipeline improve mycobiome data. PeerJ 6:4925. https://doi.org/10.7717/peerj.4925

35. Edgar RC, (2010) Search and clustering orders of magnitude faster than BLAST. Bioinformatics 26:2460–2461. https://doi.org/10.1093/bioinformatics/btq461

36. Edgar RC, Flyvbjerg H, (2015) Error filtering, pair assembly and error correction for next-generation sequencing reads. Bioinformatics 31:3476–3482. https://doi.org/10.1093/bioinformatics/btv401

37. Rognes T, Flouri T, Nichols B, Quince C, Mahé F (2016) VSEARCH: a versatile open source tool for metagenomics. PeerJ 4:2584. https://doi.org/10.7717/peerj.2584

38. Nilsson RH, Larsson KH, Taylor AFS, Bengtsson-Palme J, Jeppesen TS, Schigel D, Kennedy P, Picard K, Glöckner FO, Tedersoo L, Saar I (2018) The UNITE database for molecular identification of fungi: handling dark taxa and parallel taxonomic classifications. Nucleic acids research 47:D259–D264. https://doi.org/10.1093/nar/gky1022

39. Bolyen E et al. (2019). Reproducible, interactive, scalable and extensible microbiome data science using QIIME Nature biotechnology 37: 852–857. https://doi.org/10.1038/s41587-019-0209-9

40. Callahan BJ, McMurdie PJ, Rosen MJ, Han AW, Johnson AJA, Holmes SP (2016) DADA2: high-resolution sample inference from Illumina amplicon data. Nature methods 13:581. https://doi.org/10.1038/nmeth.3869

41. Bokulich NA, Kaehler BD, Rideout JR, Dillon M, Bolyen E, Knight R, Huttley GA Caporaso JG (2018) Optimizing taxonomic classification of marker-gene amplicon sequences with QIIME 2’s q2-feature-classifier plugin. Microbiome 6:90. https://doi.org/10.1186/s40168-018-0470-z

42. Quast C, Pruesse E, Yilmaz P, Gerken J, Schweer T, Yarza P, Peplies J, Glöckner FO, (2012) The SILVA ribosomal RNA gene database project: improved data processing and web-based tools. Nucleic acids research 41:D590–D596. https://doi.org/10.1093/nar/gks1219

43. R Core Team (2018) R: A Language and Environment for Statistical Computing. R Foundation for Statistical Computing., Vienna, Austria URL https://www.R-project.org.

44. RStudio Team (2016) RStudio: Integrated Development for R. RStudio, Inc., Boston, MA URL http://www.rstudio.com/.

45. McMurdie PJ, Holmes S (2013) phyloseq: an R package for reproducible interactive analysis and graphics of microbiome census data. PloS one 8:61217. https://doi.org/10.1371/journal.pone.0061217

46. Oksanen J, Blanchet FG, Kindt R, Legendre P, Minchin PR, O’hara RB, Simpson GL, Solymos P, Stevens MHH, Wagner H, Oksanen MJ (2019) Package ‘vegan’. Community ecology package, version, 2.5-4.

47. Kurtz ZD, Müller CL, Miraldi ER, Littman DR, Blaser MJ, Bonneau RA (2015) Sparse and compositionally robust inference of microbial ecological networks. PLoS computational biology 11:1004226. https://doi.org/10.1371/journal.pcbi.1004226

48. Tipton L, Müller CL, Kurtz ZD, Huang L, Kleerup E, Morris A, Bonneau R, Ghedin E (2018) Fungi stabilize connectivity in the lung and skin microbial ecosystems. Microbiome 6:12. https://doi.org/10.1186/s40168-017-0393-0

49. Gu Z, Gu L, Eils R, Schlesner M, Brors B (2014) circlize implements and enhances circular visualization in R. Bioinformatics 30:2811–2812. https://doi.org/10.1093/bioinformatics/btu393

50. Roberts DW (2016) labdsv: Ordination and Multivariate Analysis for Ecology. R package version 1.8-0.

51. Bowker MA, Belnap J, Büdel B, Sannier C, Pietrasiak N, Eldridge DJ, Rivera-Aguilar V. (2016) Controls on distribution patterns of biological soil crusts at micro-to global scales. In: Biological soil crusts: an organizing principle in drylands. Springer, Cham, pp 173–197

52. Büdel, B. (2001). Synopsis: comparative biogeography of soil-crust biota. In: Biological soil crusts: structure, function, and management. Springer, Berlin, Heidelberg, pp 141–152

53. Holmgren CA, Betancourt JL, Rylander K A (2010). A long□term vegetation history of the Mojave– Colorado desert ecotone at Joshua Tree National Park. Journal of Quaternary Science, 25: 222–236 https://doi.org/10.1002/jqs.1313

54. Chen Q, Jiang JR, Zhang GZ, Cai L, Crous PW (2015) Resolving the Phoma enigma. Studies in mycology 82: 137–217. https://doi.org/10.1016/j.simyco.2015.10.003

55. Tedersoo L, Bahram M, Põlme S, Kõljalg U, Yorou NS, Wijesundera R, Ruiz LV, Vasco-Palacios AM, Thu PQ, Suija A, Smith ME (2014) Global diversity and geography of soil fungi. science 346:1256688. https://doi.org/10.1126/science.1256688

56. Duncan CG, Eslyn WE (1966) Wood-decaying ascomycetes and fungi imperfecti. Mycologia 58:642–645 https://doi.org/10.2307/3757045

57. Worrall JJ, Anagnost SE, Zabel RA (1997) Comparison of wood decay among diverse lignicolous fungi. Mycologia 89:199–219. https://doi.org/10.1080/00275514.1997.12026772

58. Cannon PF, Kirk PM (2007) Fungal Families of the World. CAB. International, Wallingford, Oxfordshire, UK

59. Collins SL, Sinsabaugh RL, Crenshaw C, Green L, Porras□Alfaro A, Stursova M, Zeglin LH (2008). Pulse dynamics and microbial processes in aridland ecosystems. Journal of Ecology 96: 413–420. https://doi.org/10.1111/j.1365-2745.2008.01362.x

60. Zhang Y, Aradottir AL, Serpe M, Boeken B. (2016). Interactions of biological soil crusts with vascular plants. In: Biological soil crusts: an organizing principle in drylands. Springer, Cham, pp 385–406

61. Porras-Alfaro A, Herrera J, Natvig DO, Lipinski K, Sinsabaugh RL (2011) Diversity and distribution of soil fungal communities in a semiarid grassland. Mycologia 103:10–21. https://doi.org/10.3852/09-297

62. Chilton AM, Neilan BA, Eldridge DJ (2017) Biocrust morphology is linked to marked differences in microbial community composition. Plant and Soil 1–11. https://doi.org/10.1007/s11104-017-3442-3

63. Kidron GJ, Benenson I (2014) Biocrusts serve as biomarkers for the upper 30 cm soil water content. Journal of Hydrology 509:398–405 https://doi.org/10.1016/j.jhydrol.2013.11.041

64. Doherty KD, Antoninka AJ, Bowker MA, Ayuso SV, Johnson NC (2015) A novel approach to cultivate biocrusts for restoration and experimentation. Ecological Restoration 33:13–16.

65. Antoninka A, Bowker MA, Reed SC, Doherty K (2016) Production of greenhouse□grown biocrust mosses and associated cyanobacteria to rehabilitate dryland soil function. Restoration Ecology, 24:324–335. https://doi.org/10.1111/rec.12311

66. Chiquoine LP, Abella SR, Bowker MA (2016) Rapidly restoring biological soil crusts and ecosystem functions in a severely disturbed desert ecosystem. Ecological Applications 26:1260–1272. https://doi.org/10.1002/15-0973

67. Ayuso SV, Silva AG, Nelson C, Barger NN, Garcia-Pichel F (2017) Microbial nursery production of high-quality biological soil crust biomass for restoration of degraded dryland soils. Appl. Environ. Microbiol. 83:02179–16. https://doi.org/10.1128/AEM.02179-16

68. Bowker MA (2007) Biological soil crust rehabilitation in theory and practice: an underexploited opportunity. Restoration Ecology 15:13–23. https://doi.org/10.1111/j.1526-100X.2006.00185.x

69. Read CF, Duncan DH, Vesk PA, Elith J (2011) Surprisingly fast recovery of biological soil crusts following livestock removal in southern Australia. Journal of Vegetation Science 22:905–916. https://doi.org/10.1111/j.1654-1103.2011.01296.x

70. Garcia-Pichel F, Loza V, Marusenko Y, Mateo P, Potrafka RM (2013) Temperature drives the continental-scale distribution of key microbes in topsoil communities. Science 340:1574–1577. https://doi.org/10.1126/science.1236404

