## Supplemental Tables for "Insights into the desert living skin microbiome: geography, soil depth, and crust type affect biocrust microbial communities and networks in Mojave Desert, USA"

**TableS1: Alpha diversity comparison for each fungal taxonomic class by soil depth (t-test, P < 0.05)**

| **Fungal taxonomic classes** | **Statistical analysis** |
| --- | --- |
| Sordariomycetes | t-test, t(37.95) = 4.4304, p = 7.739e-5 |
| Agaricomycetes | t-test, t(38.85) = 3.9206, p = 0.0003484 |
| Schizosaccharomycetes | t-test, t(30.315) = 3.3587, p = 0.002126 |
| Mucoromycetes | t-test, t(29.836) = 3.896, p = 0.0005113 |
| Saccharomycetes | t-test, t(29.078) = 3.1271, p = 0.003988 |
| Orbiliomycetes | t-test, t(42.793) = 2.9179, p = 0.005595 |
| Entomophthoromycetes | t-test, t(22) = 3.0262, p = 0.006206 |
| Mortierellomycetes | t-test, t(33.471) = 3.3504, p = 0.00201 |
| Basidiobolomycetes | t-test, t(30.414) = 4.7942, p = 4.038e-5 |
| Pneumocystidomycetes | t-test, t(31.281) = 2.5878, p = 0.01452 |

**TableS2: Alpha diversity comparison for each fungal taxonomic class by crust type (ANOVA, P < 0.05)**

| **Fungal taxonomic classes** | **Statistical analysis** |
| --- | --- |
| Sordariomycetes | ANOVA, F(4,19) = 5.009, p = 0.00629 |
| Eurotiomycetes | ANOVA, F(4,19) = 5.843, p = 0.00306 |
| Lecanoromycetes | ANOVA, F(4,19) = 3.189, p = 0.0366 |
| Dothideomycetes | ANOVA, F(4,19) = 14.69, p = 1.26e-5 |
| Leotiomycetes | ANOVA, F(4,19) = 3.133, p = 0.0388 |
| Agaricomycetes | ANOVA, F(4,19) = 4.288, p = 0.0122 |
| Schizosaccharomycetes | ANOVA, F(4,19) = 3.713, p = 0.0214 |
| Pezizomycetes | ANOVA, F(4,19) = 4.347, p = 0.0116 |
| Tremellomycetes | ANOVA, F(4,19) = 4.038, p = 0.0155 |

**TableS3: Alpha diversity comparison for each bacterial phylum by site (ANOVA, P < 0.05)**

| **Bacterial phylum** | **Statistical analysis** |
| --- | --- |
| Proteobacteria | ANOVA, F(3,20) = 3.86, p = 0.025 |
| Firmicutes | ANOVA, F(3,20) = 4.718, p = 0.012 |
| Bacteroidetes | ANOVA, F(3,20) = 3.354, p = 0.0394 |
| Actinobacteria | ANOVA, F(3,20) = 6.433, p = 0.00316 |
| Acidobacteria | ANOVA, F(3,20) = 8.03, p = 0.00105 |
| Planctomycetes | ANOVA, F(3,20) = 5.36, p = 0.00713 |
| Patescibacteria | ANOVA, F(3,20) = 12.937, p = 0.00212 |
| Armatimonadetes | ANOVA, F(3,20) = 3.748, p = 0.00276 |
| Gemmatimonadetes | ANOVA, F(3,20) = 3.467, p = 0.00355 |
| Verrucomicrobia | ANOVA, F(3,20) = 5.646, p = 0.00571 |

**TableS4: Alpha diversity comparison for each bacterial phylum by soil depth (t-test, P < 0.05)**

| **Bacterial phylum** | **Statistical analysis** |
| --- | --- |
| Proteobacteria | t-test, t(36.822) = 6.3377, p = 2.239e-7 |
| Firmicute | t-test, t(36.228) = 4.2031, p = 0.0001645 |
| Actinobacteria | t-test, t(30.442) = 11.272, p = 2.165e-12 |
| Euryarchaeota | t-test, t(22.497) = 4.745, p = 9.27e-5 |
| Nanoarchaeota | t-test, t(25.008) = 2.6622, p = 0.01337 |
| Thaumarchaeota | t-test, t(27.773) = 13.6, p = 8.437e-14 |
| Acidobacteria | t-test, t(26.672) = 9.481, p = 4.956e-10 |
| Planctomycetes | t-test, t(27.949) = 11.045, p = 1.05e-11 |
| Patescibacteria | t-test, t(24.532) = 5.603, p = 8.439e-6 |
| Elusimicrobia | t-test, t(24.241) = 5.2623, p = 2.078e-5 |
| Armatimonadetes | t-test, t(26.549) = 8.6557, p = 3.287e-9 |
| Chloroflexi | t-test, t(26.877) = 10.751, p = 3.1013e-1 |
| Gemmatimonadetes | t-test, t(27.926) = 8.8107, p = 1.491e-9 |
| Entotheonellaeota | t-test, t(22) = 3.2331, p = 0.003822 |
| Cyanobacteria | t-test, t(44.804) = 3.8101, p = 0.0004205 |
| Nitrospirae | t-test, t(24.256) = 8.6182, p = 7.556e-9 |
| FBP | t-test, t(39.834) = 4.9984, p = 1.201e-5 |
| Fibrobacteres | t-test, t(32.443) = 2.2399, p = 0.03207 |
| Verrucomicrobia | t-test, t(34.441) = 4.9674, p = 1.837e-5 |

**TableS5: Alpha diversity comparison for each bacterial phylum by crust type (ANOVA, P < 0.05)**

| **Bacterial phylum** | **Statistical analysis** |
| --- | --- |
| Proteobacteria | ANOVA, F(4,19) = 7.303, p = 0.000972 |
| Acidobacteria | ANOVA, F(4,19) = 4.687, p = 0.00841 |
| Planctomycetes | ANOVA, F(4,19) = 3.426, p = 0.0286 |
| Patescibacteria | ANOVA, F(4,19) = 7.917, p = 0.0292 |
| Armatimonadetes | ANOVA, F(4,19) = 2.92, p = 0.0487 |
| Deinococcus-Thermus | ANOVA, F(4,19) = 4.591, p = 0.00919 |
| Chloroflexi | ANOVA, F(4,19) = 3.564, p = 0.0249 |
| Cyanobacteria | ANOVA, F(4,19) = 10.54, p = 0.000113 |
| FBP | ANOVA, F(4,19) = 3.929, p = 0.0173 |
| Verrucomicrobia | ANOVA, F(4,19) = 5.511, p = 0.00405 |

**TableS6: Fungal indicator species by crust type**

| **Crust_Type** | **Phylum** | **Class** | **Order** | **Family** | **Genus** | **Species** |
| --- | --- | --- | --- | --- | --- | --- |
| SMC | Ascomycota | Sordariomycetes | Sordariales | Chaetomiaceae | Acrophialophora | Acrophialophora levis |
| RMC | Ascomycota | Dothideomycetes | Pleosporales | Sporormiaceae | Sporormia | Sporormia subticinensis |

**TableS7: Fungal indicator species by site**

| **Site** | **Phylum** | **Class** | **Order** | **Family** | **Genus** | **Species** |
| --- | --- | --- | --- | --- | --- | --- |
| CIMA | Ascomycota | Dothideomycetes | NA | NA | Catenulomyces | Catenulomyces convolutus |
|  | Ascomycota | Dothideomycetes | Pleosporales | Sporormiaceae | Preussia | Preussia terricola |
| JTNP | Ascomycota | Dothideomycetes | Pleosporales | Pleosporaceae | Curvularia | Curvularia inaequalis |
|  | Basidiomycota | Agaricomycetes | Agaricales | Entolomataceae | Entoloma | Entoloma halophilum |
|  | Ascomycota | Dothideomycetes | Pleosporales | Sporormiaceae | Preussia | Preussia africana |
|  | Ascomycota | Dothideomycetes | Pleosporales | Didymellaceae | Allophoma | Allophoma labilis |
| KELSO | Ascomycota | Sordariomycetes | Hypocreales | Nectriaceae | Fusarium | Fusarium oxysporum |
|  | Ascomycota | Dothideomycetes | Capnodiales | Cladosporiaceae | Cladosporium | Cladosporium herbarum |
|  | Ascomycota | Dothideomycetes | Pleosporales | Pleosporaceae | Alternaria | Alternaria hungarica |
|  | Ascomycota | Sordariomycetes | Glomerellales | Glomerellaceae | Colletotrichum | Colletotrichum gloeosporioides |
|  | Ascomycota | Dothideomycetes | Pleosporales | Pleosporaceae | Ulocladium | Ulocladium dauci |

**TableS8: Bacterial indicator species by crust type**

| **Crust type** | **Phylum** | **Class** | **Order** | **Family** | **Genus** | **Species** |
| --- | --- | --- | --- | --- | --- | --- |
| CLC | Armatimonadetes | uncultured | metagenome | metagenome | metagenome | metagenome |
|  | Bacteroidetes | Bacteroidia | Cytophagales | Spirosomaceae | Larkinella | uncultured bacterium |
|  | Bacteroidetes | Bacteroidia | Cytophagales | Microscillaceae | Flexibacter | uncultured Bacteroidetes bacterium |
|  | Chloroflexi | Chloroflexia | Kallotenuales | AKIW781 | uncultured bacterium | uncultured bacterium |
|  | Cyanobacteria | Oxyphotobacteria | Nostocales | uncultured | uncultured bacterium | uncultured bacterium |
|  | Chloroflexi | Anaerolineae | SBR1031 | A4b | metagenome | metagenome |
|  | Proteobacteria | Alphaproteobacteria | Azospirillales | Azospirillaceae | Azospirillum | Azospirillum soli |
|  | Chloroflexi | Chloroflexia | Kallotenuales | AKIW781 | uncultured Chloroflexi bacterium | uncultured Chloroflexi bacterium |
|  | Chloroflexi | Chloroflexia | Chloroflexales | Chloroflexaceae | FFCH7168 | uncultured bacterium |
| LAC | Bacteroidetes | Bacteroidia | Cytophagales | Spirosomaceae | uncultured | uncultured Bacteroidetes bacterium |
|  | Proteobacteria | Alphaproteobacteria | Acetobacterales | Acetobacteraceae | Belnapia | Belnapia moabensis |
|  | Acidobacteria | Blastocatellia (Subgroup 4) | Pyrinomonadales | Pyrinomonadaceae | RB41 | uncultured Acidobacterium sp. |
|  | Proteobacteria | Alphaproteobacteria | Acetobacterales | Acetobacteraceae | uncultured | uncultured Paracraurococcus sp. |
| RMC | Acidobacteria | Blastocatellia (Subgroup 4) | Pyrinomonadales | Pyrinomonadaceae | RB41 | uncultured Acidobacterium sp. |
|  | Chloroflexi | Chloroflexia | Chloroflexales | Chloroflexaceae | Chloronema | uncultured bacterium |
|  | Proteobacteria | Alphaproteobacteria | Rhizobiales | Beijerinckiaceae | uncultured | Salinarimonas sp. BN140002 |
|  | FBP | uncultured bacterium | uncultured bacterium | uncultured bacterium | uncultured bacterium | uncultured bacterium |
|  | Bacteroidetes | Bacteroidia | Cytophagales | Cytophagaceae | Rhodocytophaga | uncultured bacterium |
|  | Bacteroidetes | Bacteroidia | Cytophagales | Cyclobacteriaceae | uncultured | uncultured Bacteroidetes bacterium |
| SMC | Deinococcus-Thermus | Deinococci | Deinococcales | Deinococcaceae | Deinococcus | Deinococcus pimensis DSM 21231 |
|  | Planctomycetes | Phycisphaerae | Tepidisphaerales | WD2101 soil group | uncultured planctomycete | uncultured planctomycete |
|  | Cyanobacteria | Oxyphotobacteria | Nostocales | Nostocaceae | Calothrix PCC-6303 | Calothrix sp. HA4186-MV5 |
|  | Proteobacteria | Alphaproteobacteria | Caulobacterales | Caulobacteraceae | PMMR1 | uncultured bacterium |
|  | Cyanobacteria | Oxyphotobacteria | Nostocales | uncultured | uncultured bacterium | uncultured bacterium |
|  | Planctomycetes | Planctomycetacia | Gemmatales | Gemmataceae | uncultured | uncultured bacterium |
|  | FBP | uncultured bacterium | uncultured bacterium | uncultured bacterium | uncultured bacterium | uncultured bacterium |
|  | Chloroflexi | Chloroflexia | Kallotenuales | AKIW781 | uncultured bacterium | uncultured bacterium |
|  | Acidobacteria | Blastocatellia (Subgroup 4) | Blastocatellales | Blastocatellaceae | uncultured | uncultured bacterium |
|  | Proteobacteria | Alphaproteobacteria | Acetobacterales | Acetobacteraceae | Roseomonas | uncultured bacterium |
|  | Proteobacteria | Alphaproteobacteria | Rhizobiales | Beijerinckiaceae | Psychroglaciecola | uncultured bacterium |
|  | Actinobacteria | Actinobacteria | Frankiales | Sporichthyaceae | uncultured | uncultured bacterium |
|  | Cyanobacteria | Oxyphotobacteria | Nostocales | Chroococcidiopsaceae | uncultured | uncultured bacterium |
|  | Planctomycetes | Planctomycetacia | Isosphaerales | Isosphaeraceae | Singulisphaera | uncultured bacterium |
|  | Proteobacteria | Alphaproteobacteria | Acetobacterales | Acetobacteraceae | Craurococcus | uncultured bacterium |
|  | Planctomycetes | Phycisphaerae | Tepidisphaerales | WD2101 soil group | uncultured planctomycete | uncultured planctomycete |
|  | Acidobacteria | Blastocatellia (Subgroup 4) | Blastocatellales | Blastocatellaceae | Blastocatella | uncultured bacterium |

**TableS9: Bacterial indicator species by site**

| Site | Phylum | Class | Order | Family | Genus | Species |
| --- | --- | --- | --- | --- | --- | --- |
| CIMA | Armatimonadetes | Armatimonadia | Armatimonadales | uncultured bacterium | uncultured bacterium | uncultured bacterium |
|  | Gemmatimonadetes | Longimicrobia | Longimicrobiales | Longimicrobiaceae | uncultured bacterium | uncultured bacterium |
|  | Proteobacteria | Alphaproteobacteria | Acetobacterales | Acetobacteraceae | Craurococcus | uncultured proteobacterium |
|  | Armatimonadetes | Armatimonadia | Armatimonadales | uncultured soil bacterium | uncultured soil bacterium | uncultured soil bacterium |
|  | Bacteroidetes | Bacteroidia | Cytophagales | Spirosomaceae | uncultured | uncultured bacterium |
|  | Proteobacteria | Alphaproteobacteria | Caulobacterales | Caulobacteraceae | PMMR1 | uncultured bacterium |
|  | Chloroflexi | Chloroflexia | Kallotenuales | AKIW781 | uncultured bacterium | uncultured bacterium |
|  | Acidobacteria | Acidobacteriia | Solibacterales | Solibacteraceae (Subgroup 3) | Bryobacter | uncultured Acidobacteria bacterium |
|  | Armatimonadetes | Armatimonadia | Armatimonadales | uncultured bacterium | uncultured bacterium | uncultured bacterium |
|  | Chloroflexi | Chloroflexia | Chloroflexales | Herpetosiphonaceae | Herpetosiphon | uncultured bacterium |
|  | Bacteroidetes | Bacteroidia | Chitinophagales | Chitinophagaceae | Segetibacter | Segetibacter aerophilus |
|  | Armatimonadetes | Armatimonadia | Armatimonadales | uncultured bacterium | uncultured bacterium | uncultured bacterium |
|  | Bacteroidetes | Bacteroidia | Cytophagales | Hymenobacteraceae | Nibribacter | uncultured bacterium |
|  | FBP | uncultured soil bacterium | uncultured soil bacterium | uncultured soil bacterium | uncultured soil bacterium | uncultured soil bacterium |
|  | Planctomycetes | Planctomycetacia | Isosphaerales | Isosphaeraceae | uncultured | uncultured Isosphaera sp. |
|  | Cyanobacteria | Oxyphotobacteria | Nostocales | Chroococcidiopsaceae | Chroococcidiopsis SAG 2023 | Chroococcidiopsis sp. BB79.2 |
|  | Cyanobacteria | Oxyphotobacteria | Nostocales | uncultured | uncultured bacterium | uncultured bacterium |
|  | Cyanobacteria | Oxyphotobacteria | Nostocales | Chroococcidiopsaceae | Chroococcidiopsis SAG 2023 | uncultured bacterium |
|  | Chloroflexi | Chloroflexia | Chloroflexales | Roseiflexaceae | uncultured | uncultured Chloroflexi bacterium |
|  | Bacteroidetes | Bacteroidia | Cytophagales | Spirosomaceae | uncultured | uncultured Bacteroidetes bacterium |
|  | Cyanobacteria | Oxyphotobacteria | Nostocales | uncultured | uncultured bacterium | uncultured bacterium |
|  | Deinococcus-Thermus | Deinococci | Deinococcales | Trueperaceae | Truepera | uncultured endolithic bacterium |
|  | Bacteroidetes | Bacteroidia | Cytophagales | Cytophagaceae | uncultured | uncultured bacterium |
|  | Chloroflexi | Chloroflexia | Thermomicrobiales | JG30-KF-CM45 | uncultured soil bacterium | uncultured soil bacterium |
|  | Verrucomicrobia | Verrucomicrobiae | Chthoniobacterales | Chthoniobacteraceae | Chthoniobacter | uncultured bacterium |
|  | Cyanobacteria | Oxyphotobacteria | Nostocales | uncultured | uncultured bacterium | uncultured bacterium |
|  | Bacteroidetes | Bacteroidia | Chitinophagales | uncultured | metagenome | metagenome |
|  | Deinococcus-Thermus | Deinococci | Deinococcales | Deinococcaceae | Deinococcus | Deinococcus maricopensis DSM 21211 |
|  | Cyanobacteria | Oxyphotobacteria | Nostocales | Chroococcidiopsaceae | Chroococcidiopsis SAG 2023 | Chroococcidiopsis sp. BB79.2 |
| GMT | Cyanobacteria | Oxyphotobacteria | Nostocales | uncultured | uncultured cyanobacterium | uncultured cyanobacterium |
|  | Cyanobacteria | Oxyphotobacteria | Nostocales | Nostocaceae | uncultured | uncultured bacterium |
|  | Chloroflexi | TK10 | uncultured bacterium | uncultured bacterium | uncultured bacterium | uncultured bacterium |
|  | Cyanobacteria | Oxyphotobacteria | Nostocales | Nostocaceae | Mastigocladopsis PCC-10914 | uncultured bacterium |
|  | Planctomycetes | Phycisphaerae | Tepidisphaerales | WD2101 soil group | uncultured planctomycete | uncultured planctomycete |
|  | Bacteroidetes | Bacteroidia | Cytophagales | Hymenobacteraceae | Hymenobacter | Parahymenobacter deserti |
|  | Cyanobacteria | Oxyphotobacteria | Nostocales | Nostocaceae | uncultured | uncultured bacterium |
|  | Chloroflexi | Chloroflexia | Kallotenuales | AKIW781 | uncultured bacterium | uncultured bacterium |
|  | Cyanobacteria | Oxyphotobacteria | Nostocales | Nostocaceae | Mastigocladopsis PCC-10914 | uncultured bacterium |
|  | Bacteroidetes | Bacteroidia | Cytophagales | Spirosomaceae | uncultured | uncultured Bacteroidetes bacterium |
|  | Bacteroidetes | Bacteroidia | Cytophagales | Hymenobacteraceae | Adhaeribacter | uncultured bacterium |
|  | Chloroflexi | Chloroflexia | Kallotenuales | AKIW781 | uncultured bacterium | uncultured bacterium |
|  | Armatimonadetes | uncultured | uncultured bacterium | uncultured bacterium | uncultured bacterium | uncultured bacterium |
|  | Chloroflexi | Chloroflexia | Chloroflexales | Chloroflexaceae | FFCH7168 | uncultured bacterium |
|  | Armatimonadetes | uncultured | uncultured bacterium | uncultured bacterium | uncultured bacterium | uncultured bacterium |
| JTNP | Bacteroidetes | Bacteroidia | Cytophagales | Hymenobacteraceae | Hymenobacter | Hymenobacter rigui |
|  | Gemmatimonadetes | Gemmatimonadetes | Gemmatimonadales | Gemmatimonadaceae | Gemmatimonas | metagenome |
|  | Proteobacteria | Alphaproteobacteria | Rhodobacterales | Rhodobacteraceae | Rubellimicrobium | uncultured bacterium |
|  | Bacteroidetes | Bacteroidia | Chitinophagales | Chitinophagaceae | Flavisolibacter | uncultured bacterium |
|  | Proteobacteria | Deltaproteobacteria | Myxococcales | Haliangiaceae | Haliangium | uncultured delta proteobacterium |
|  | Proteobacteria | Alphaproteobacteria | Rhodobacterales | Rhodobacteraceae | Rubellimicrobium | uncultured bacterium |
|  | Bacteroidetes | Bacteroidia | Chitinophagales | Chitinophagaceae | Cnuella | uncultured bacterium |
|  | Verrucomicrobia | Verrucomicrobiae | Pedosphaerales | Pedosphaeraceae | Pedosphaera | uncultured bacterium |
|  | Armatimonadetes | Fimbriimonadia | Fimbriimonadales | Fimbriimonadaceae | metagenome | metagenome |
|  | Verrucomicrobia | Verrucomicrobiae | Chthoniobacterales | Chthoniobacteraceae | Chthoniobacter | uncultured bacterium |
|  | Actinobacteria | Actinobacteria | Frankiales | Geodermatophilaceae | Geodermatophilus | uncultured bacterium |
|  | Bacteroidetes | Bacteroidia | Cytophagales | Spirosomaceae | Rhabdobacter | uncultured bacterium |
| KELSO | Chloroflexi | Chloroflexia | Kallotenuales | AKIW781 | uncultured bacterium | uncultured bacterium |
|  | Actinobacteria | Thermoleophilia | Solirubrobacterales | 67-14 | uncultured actinobacterium | uncultured actinobacterium |
|  | Proteobacteria | Alphaproteobacteria | Rhodobacterales | Rhodobacteraceae | Rubellimicrobium | uncultured bacterium |
|  | Proteobacteria | Alphaproteobacteria | Acetobacterales | Acetobacteraceae | Roseomonas | Roseomonas pecuniae |
|  | Proteobacteria | Alphaproteobacteria | Azospirillales | Azospirillaceae | Skermanella | uncultured bacterium |
|  | Bacteroidetes | Bacteroidia | Chitinophagales | uncultured | metagenome | metagenome |
|  | Cyanobacteria | Oxyphotobacteria | Nostocales | Coleofasciculaceae | uncultured | uncultured cyanobacterium |
|  | Proteobacteria | Deltaproteobacteria | Myxococcales | BIrii41 | uncultured bacterium | uncultured bacterium |
|  | Bacteroidetes | Bacteroidia | Cytophagales | Cytophagaceae | Rhodocytophaga | uncultured Bacteroidetes bacterium |
|  | Proteobacteria | Alphaproteobacteria | Sphingomonadales | Sphingomonadaceae | Sphingomonas | Sphingomonas kaistensis |
|  | Proteobacteria | Alphaproteobacteria | Acetobacterales | Acetobacteraceae | uncultured | uncultured Paracraurococcus sp. |
