## Supplementary figures and images for "Insights into the desert living skin microbiome: geography, soil depth, and crust type affect biocrust microbial communities and networks in Mojave Desert, USA"

### FigureS1

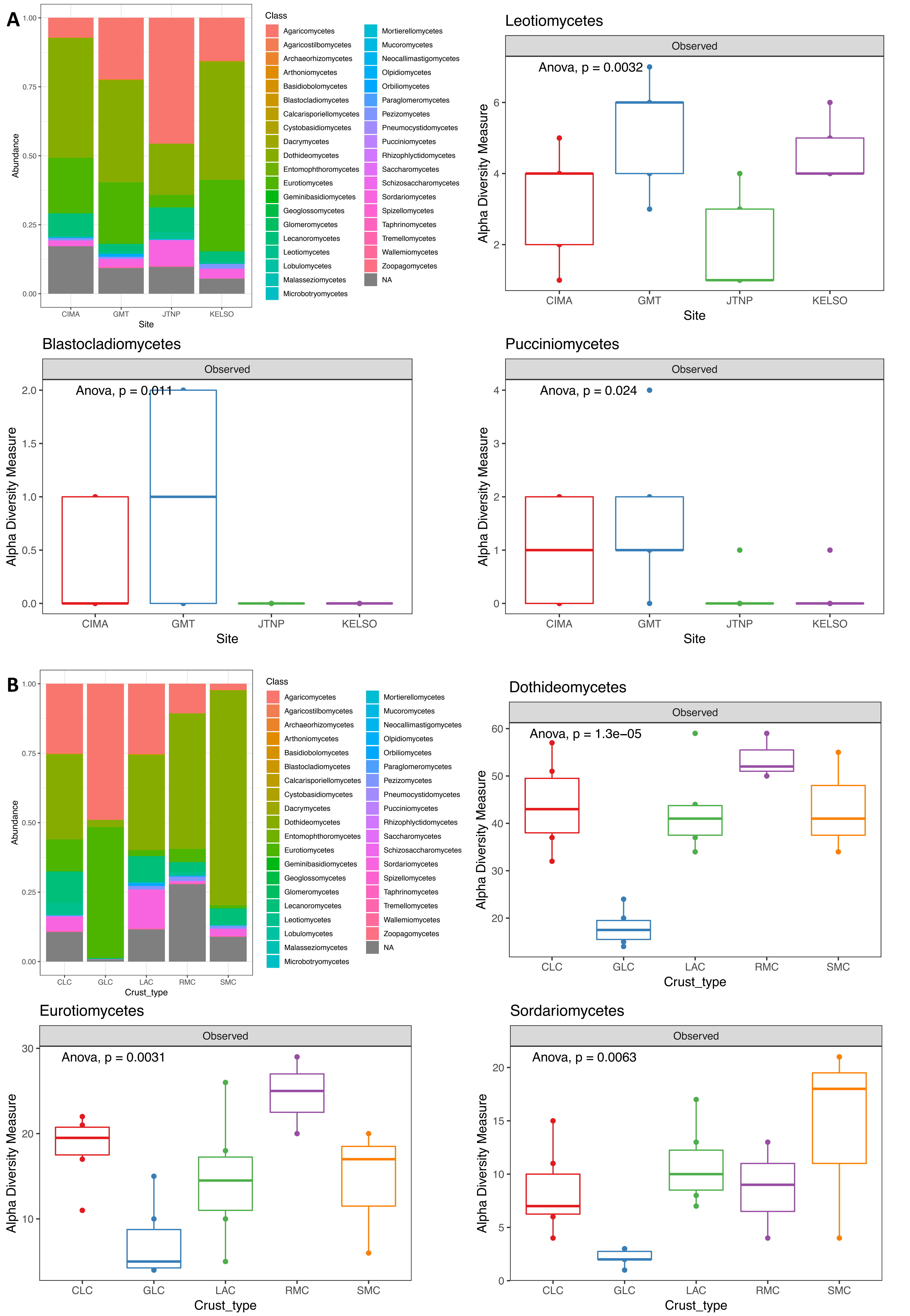

### FigureS2

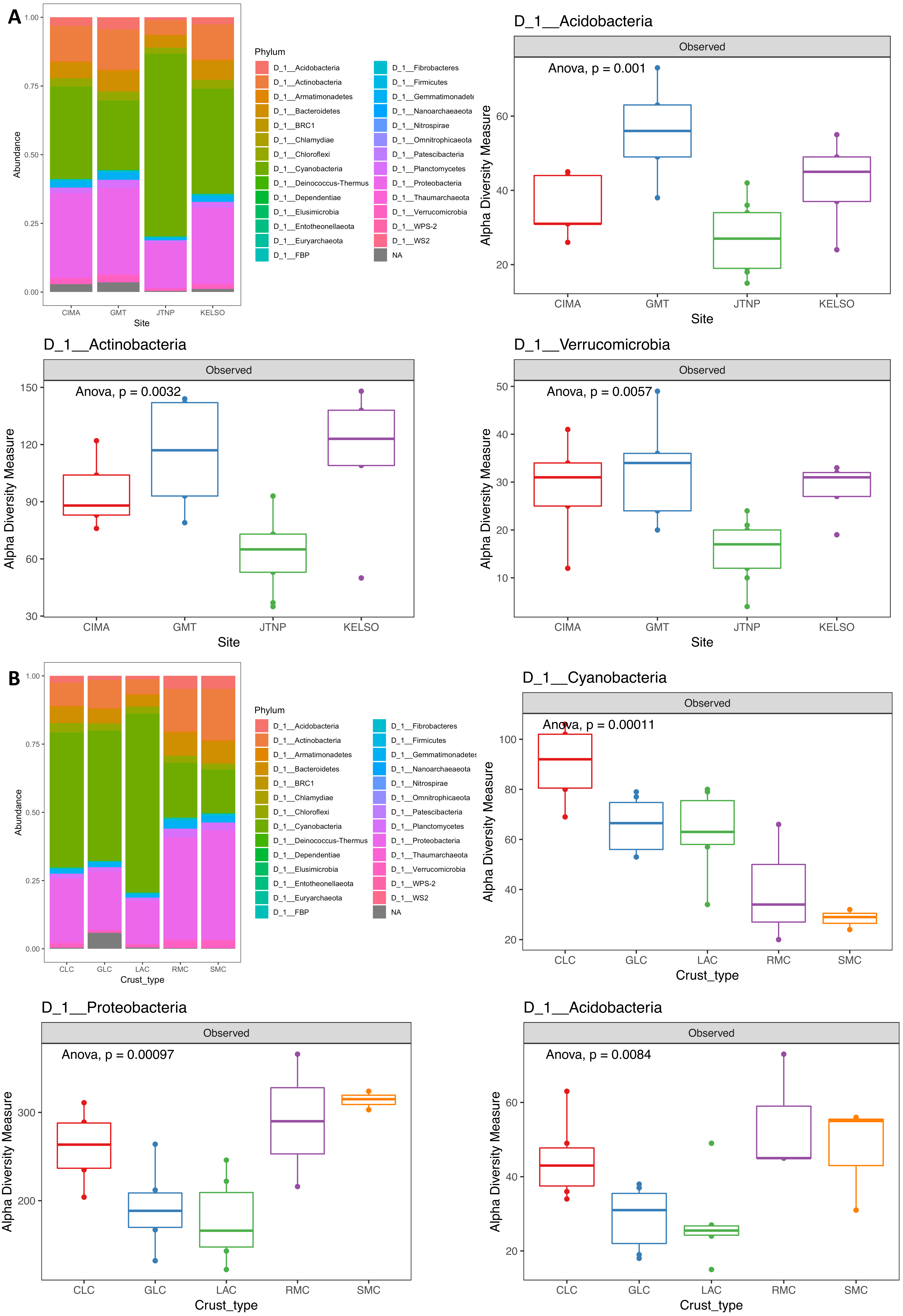

### FigureS3

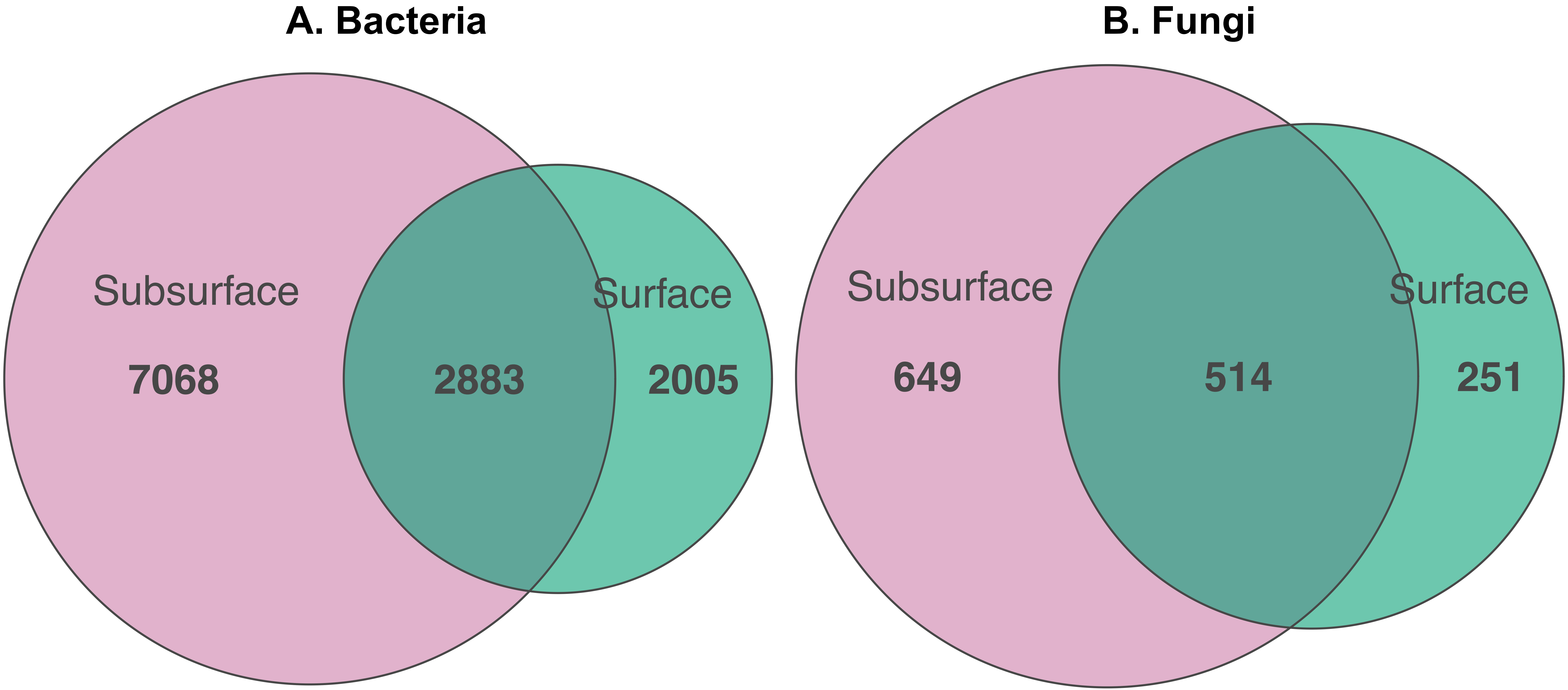

### FigureS4

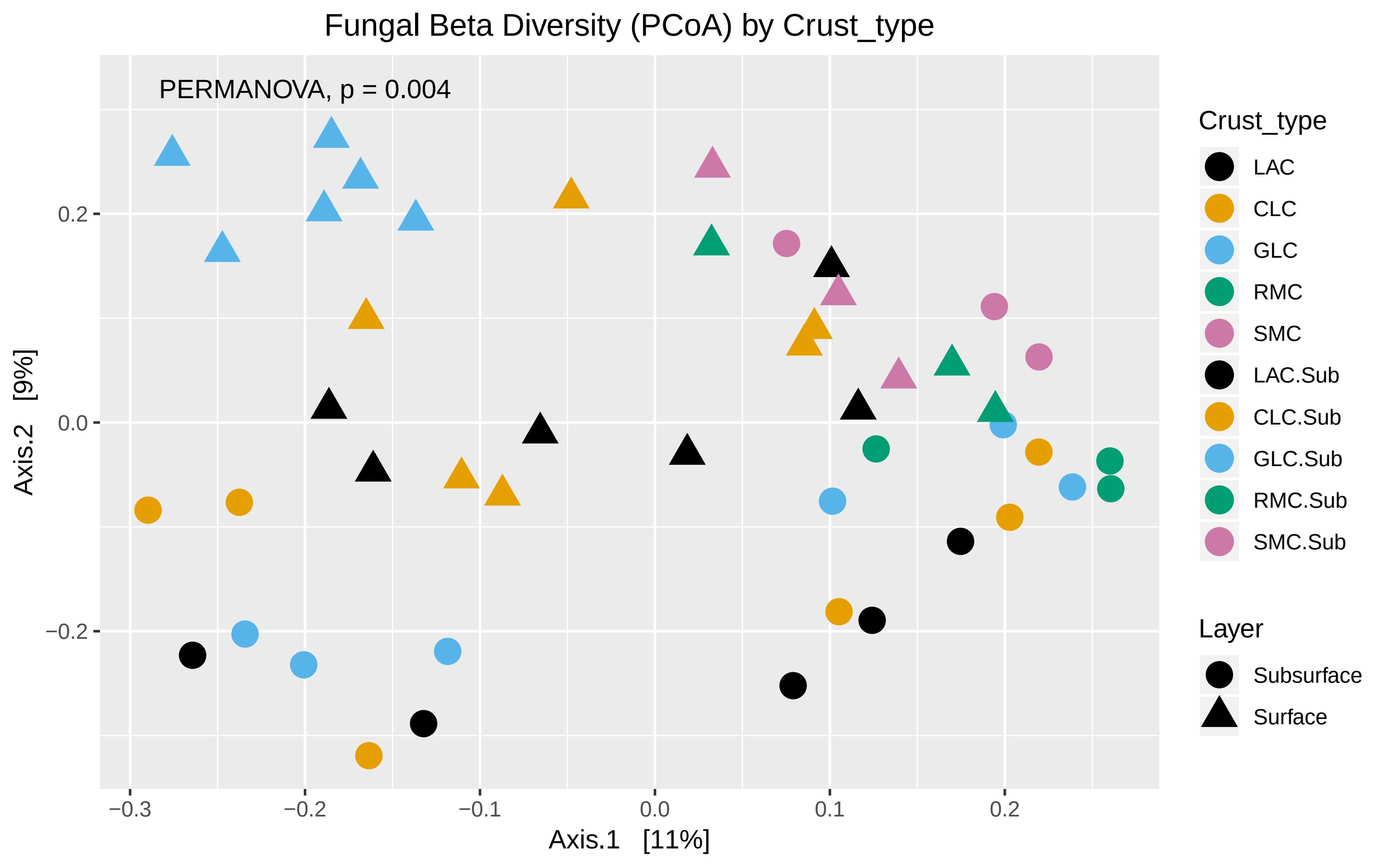

### FigureS5

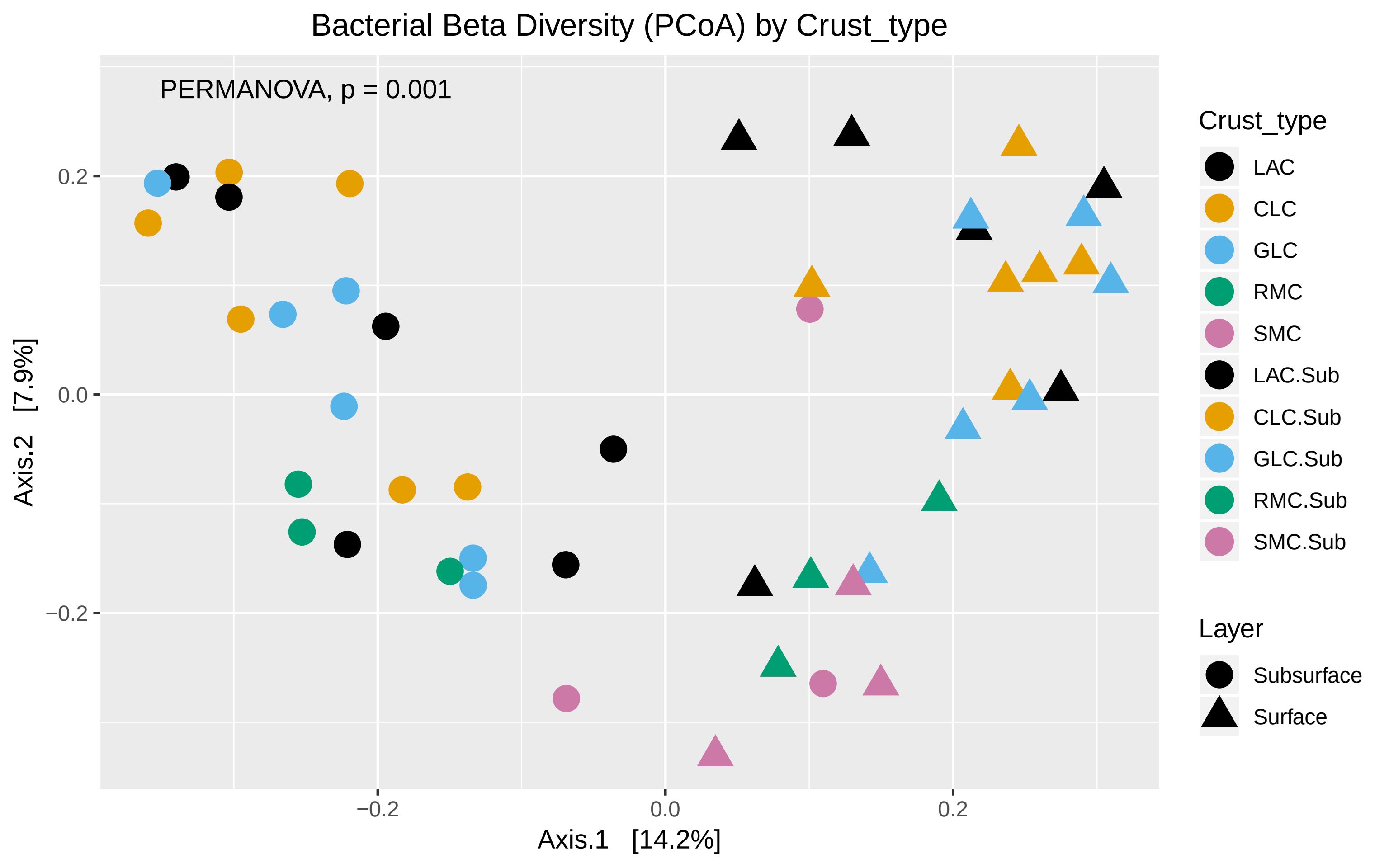

### FigureS6

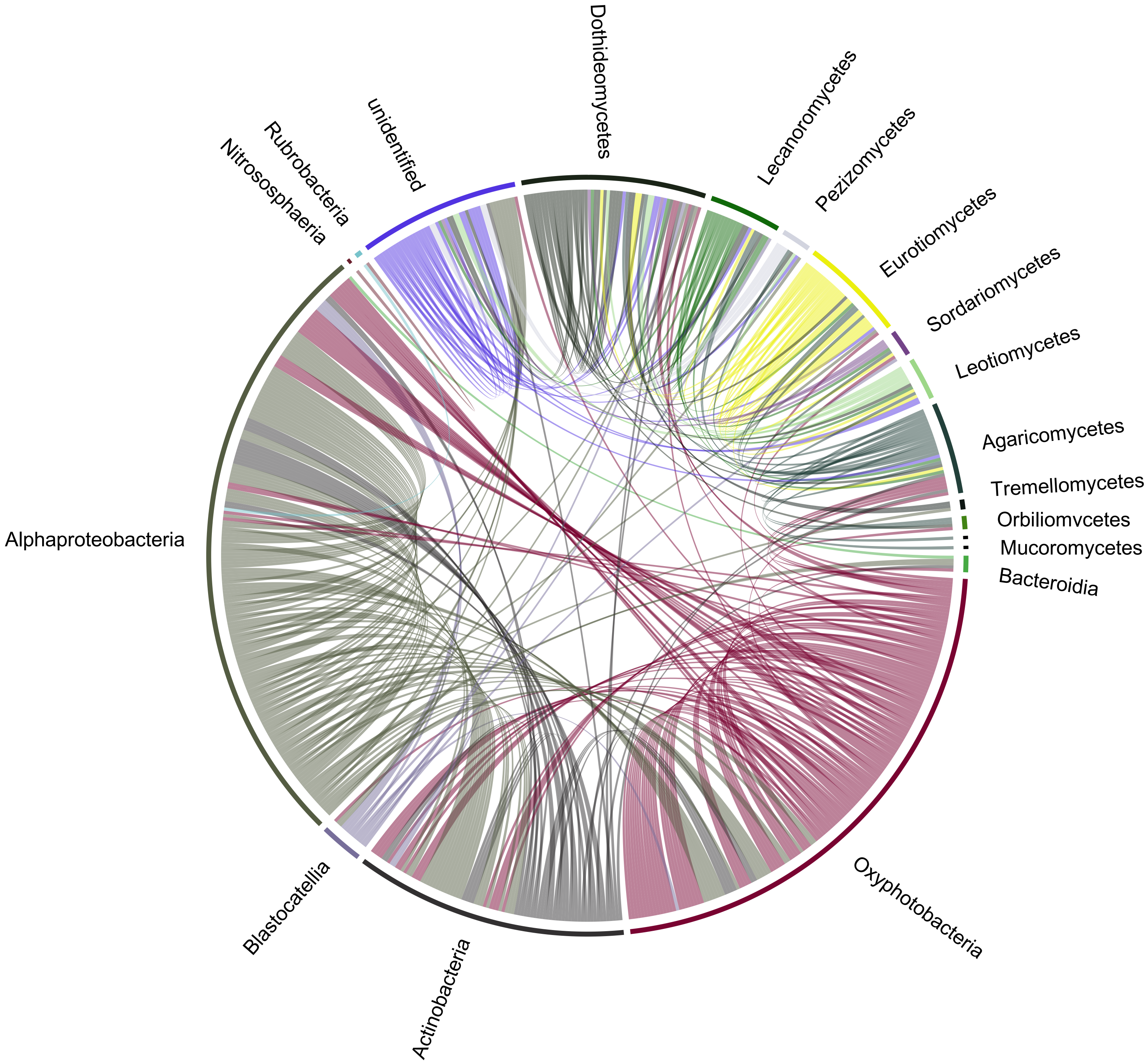

### FigureS7

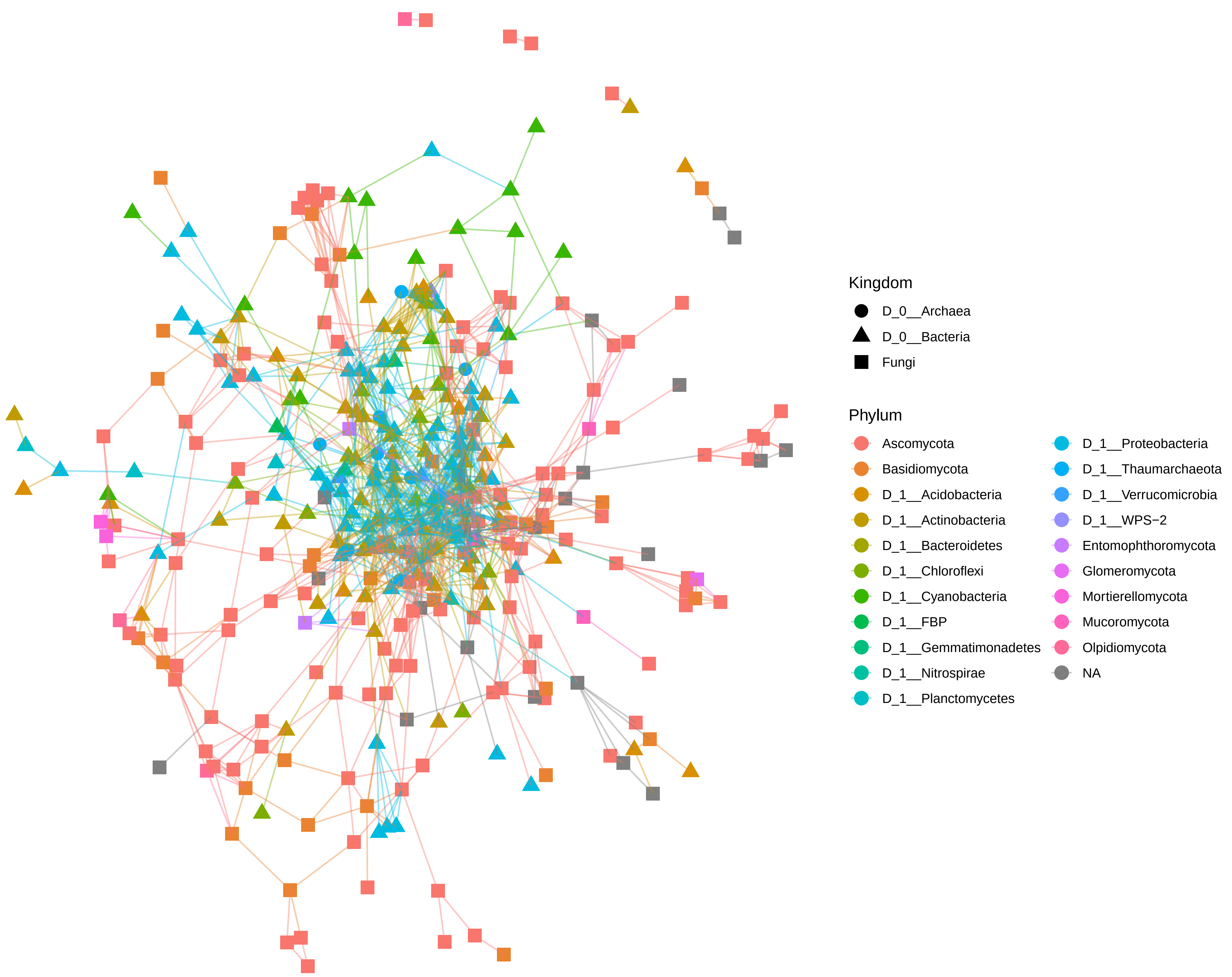

### FigureS8

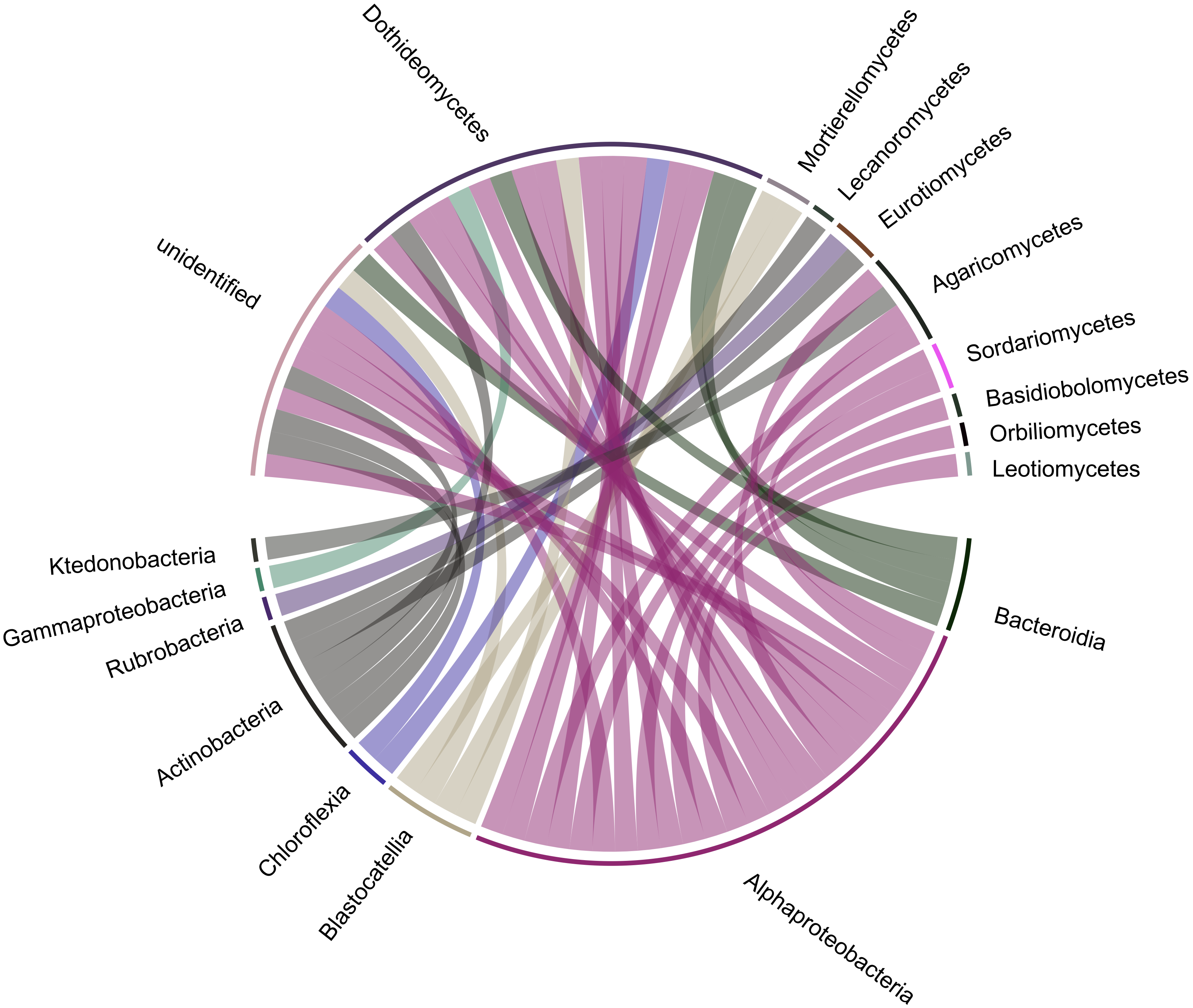
